# Single cell transcriptional analysis of human adenoids identifies molecular features of airway microfold cells

**DOI:** 10.1101/2024.10.19.619143

**Authors:** Samuel Alvarez-Arguedas, Khadijah Mazhar, Andi Wangzhou, Ishwarya Sankaranarayanan, Gabriela Gaona, John T. Lafin, Ron B. Mitchell, Theodore J. Price, Michael U. Shiloh

**Affiliations:** Department of Internal Medicine, University of Texas Southwestern Medical Center, Dallas, Texas 75390, USA; School of Behavioral and Brain Sciences, University of Texas at Dallas, Richardson, TX, 75080, USA; Center for Advanced Pain Studies, University of Texas at Dallas, Richardson, TX, 75080, USA; Department of Urology, University Texas Southwestern Medical Center, Dallas, TX 75390, USA; Department of Otolaryngology, University of Texas Southwestern Medical Center, Dallas, TX, 75390, USA; Department of Microbiology, University of Texas Southwestern Medical Center, Dallas, Texas 75390, USA

## Abstract

The nasal, oropharyngeal, and bronchial mucosa are primary contact points for airborne pathogens like *Mycobacterium tuberculosis* (Mtb), SARS-CoV-2, and influenza virus. While mucosal surfaces can function as both entry points and barriers to infection, mucosa-associated lymphoid tissues (MALT) facilitate early immune responses to mucosal antigens. MALT contains a variety of specialized epithelial cells, including a rare cell type called a microfold cell (M cell) that functions to transport apical antigens to basolateral antigen-presenting cells, a crucial step in the initiation of mucosal immunity. M cells have been extensively characterized in the gastrointestinal (GI) tract in murine and human models. However, the precise development and functions of human airway M cells is unknown. Here, using single-nucleus RNA sequencing (snRNA-seq), we generated an atlas of cells from the human adenoid and identified 16 unique cell types representing basal, club, hillock, and hematopoietic lineages, defined their developmental trajectories, and determined cell-cell relationships. Using trajectory analysis, we found that human airway M cells develop from progenitor club cells and express a gene signature distinct from intestinal M cells. Surprisingly, we also identified a heretofore unknown epithelial cell type demonstrating a robust interferon-stimulated gene signature. Our analysis of human adenoid cells enhances our understanding of mucosal immune responses and the role of M cells in airway immunity. This work also provides a resource for understanding early interactions of pathogens with airway mucosa and a platform for development of mucosal vaccines.

## Introduction

The respiratory mucosa can function both as a portal of entry and barrier to infection ^1^ for a number of endemic and epidemic pathogens such as *Mycobacterium tuberculosis* ^2,3^, SARS-CoV2 ^4,5^, influenza virus ^6^, measles virus ^7^, *Neisseria meningitidis* ^8^, and *Streptococcus pneumoniae* ^9^. In mammals, mucosal immune structures that include innate and adaptive immune cells as well as specialized epithelial cells called mucosa-associated lymphoid tissue (MALT) facilitate the development of early immune responses against inhaled pathogens. In the upper airway, MALT structures include the adenoids or pharyngeal tonsil, tubal tonsils, palatine tonsils, and lingual tonsils of Waldeyer’s ring ^10^. Finally, both bronchus-associated lymphoid tissue (BALT) and inducible BALT (iBALT) that can form in response to airway infection or inflammation line the distal bronchial airways ^11,12^.

In addition to the airway, MALT is prominent within the gastrointestinal (GI) tract, and in the GI tract is best characterized by small-intestinal Peyer’s patches ^13^. Overlying MALT in the airway and Peyer’s patches in the GI tract is a specialized cell called a microfold cell (M cell) that captures antigens on the apical surface for transcytosis to a basolateral pocket occupied by antigen-presenting immune cells ^14,15^. While GI tract M cells have been extensively studied ^15,16^, recent work has highlighted the importance of medium and lower airway M cells ^17–19^. However, neither the basic features of upper airway M cells, nor their role in the development of mucosal immunity against airway pathogens is well understood.

We and others have demonstrated that M cells are a portal of entry for Mtb in the nasal-associated lymphoid tissue (NALT) and airway ^2,3^. Mtb infection of M cells in mice and humans requires the Mtb secreted effector EsxA and a cell surface receptor on M cells, scavenger receptor B1 (SCARB1)^20^. Notably, airway infection, potentially mediated by M cells, may also explain extrapulmonary forms of tuberculosis such as scrofula and mediastinal tuberculosis ^21^.

A deeper insight into airway M cell biology and the role of M cells in initiating airway mucosal immunity is critical for our understanding of the earliest interactions of Mtb and other airway pathogens with the airway mucosa ^22^. Using single-nucleus RNA sequencing (snRNA-seq) of primary human adenoid tissue, we annotated and characterized the cell composition of the human adenoid. We determined the transcriptional profile and developmental trajectory not only of primary human M cells in the upper airway, but also identified a unique cell type that has not previously been described. This study paves the way to fully understand human mucosal immune responses to airway pathogens and the early role of airway M cells in initiating this immune response, and may also influence future mucosal vaccine strategies ^23^.

## Results

### Cell composition of the human adenoid

To decipher the cellular composition of upper airway mucosa-associated lymphoid tissue, we collected and sequenced 16,779 nuclei from 6 different human adenoids by snRNA-seq (**Figure 1A**). To visualize cell clusters in the human adenoid, we performed UMAP reduction (dimensionality = 14, resolution = 1.2), and identified 7 different nucleus populations by manually checking the expression of cell type gene expression markers for each nucleus population (**Figure 1B**, **Supplemental Figures 1A, B**). The sample composition of each identified cluster, the number of nuclei per sample, the median number of RNA molecules detected within a nucleus (median_nCount), and the median number of genes detected in each nucleus (median_nFeat) represented the technical and biological variability in the sample preparation for each human donor (**Supplemental Figures 2A, B**).

**Figure 1.**
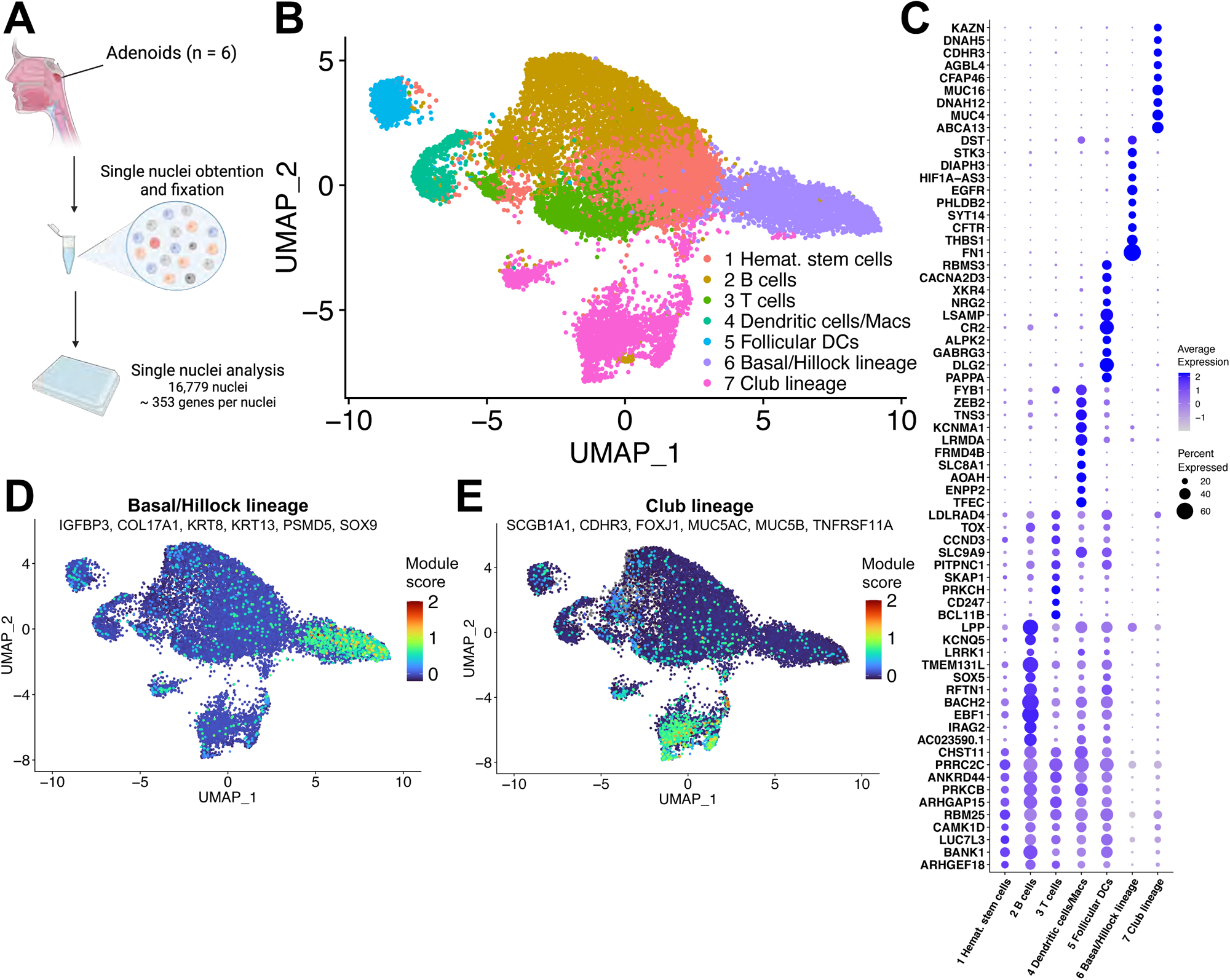
Cell composition of the human adenoid. (A) Overview of the sample collection, preparation and sequencing analysis. (B) UMAP plot of 16,779 nuclei from analyzed human adenoids (dim. = 14, res. = 1.2). (C) Dotplot of the average expression of top DEGs for each nucleus population. Module score of a list of genes associated with basal/hillock lineage (D) or club lineage (E) on the UMAP plot of the human adenoids. The module score values represent the expression values normalized to the maximum expression of the combination of genes across all the nuclei.

From 7 cell populations, we assigned 5 clusters to adaptive and innate immune cell lineages, and 2 clusters to epithelial cell lineages based on the presence or absence of protein tyrosine phosphatase receptor type C (*PTPRC*, also known as *CD45*) (**Supplemental Figures 1A, B**). The total number of nuclei isolated and sequenced representing the immune-related clusters (11,514 nuclei) was significantly greater than the epithelial clusters (5,265 nuclei) (**Supplemental Figure 2C**). To define the individual features of each population, we analyzed the number of genes detected, the total number of RNA molecules, and the percentage of mitochondrial RNA in each population (**Supplemental Figures 2C, D**). Notably, club lineage cells had a higher content of mitochondrial RNA transcripts compared to the other adenoid cell clusters (**Supplemental Figure 2D**). These data are consistent with prior observations in both the GI tract and airway that club cells, also named Clara or secretory cells, and other club lineage cells have a higher mitochondrial content and are more metabolically active in part due to highly active secretory functions ^24–26^.

The first unexpected finding from our analysis was a cluster of cells with features consistent with a hematopoietic stem cell (HSC) lineage, which, to our knowledge, have not previously been observed in the adenoid (**Supplemental Figures 1A, B**). Genes characteristic of HSC included *PTPRC*, ataxin 1 (*ATXN1*), cluster of differentiation 38 (*CD38*), and signaling lymphocytic activation molecule family member 1 (*SLAMF1*) (**Supplemental Figures 1A, B**) ^27–30^. Furthermore, the top differentially expressed genes (DEGs) for the HSC cluster were also significantly expressed in the other immune-related clusters despite their disparate final differentiation status (**Figure 1C**). This overlap identified the HSC cluster as a possible early common progenitor for all terminally differentiated immune cell clusters. A second surprising finding from the identification of potential HSC in the adenoid was that HSC genes overlapped with the transcriptional signature of follicular dendritic cells (DCs) (**Figure 1C**, **Supplemental Figure 1B**). Though previous work has identified a mesenchymal origin for follicular DCs ^31–33^, the earliest precursor of follicular DCs is not well defined. When we analyzed the other immune cell clusters, we found cell marker genes and DEGs that clearly identified each unique immune cell type (**Figure 1C**, **Supplemental Figures 1A, B**). For example, CD19 molecule (*CD19*), PX domain containing serine/threonine kinase like (*PXK*), CD74 molecule (*CD74*), the transcription factors SRY-box transcription factor 5 (*SOX5*), and EBF transcription factor 1 (*EBF1*) are associated with B cell maturation ^34-38^; *TRBC2* is the T cell receptor beta constant 2, and *CD247* and *CD3E* are the zeta chain and epsilon subunit of the CD3 T cell receptor respectively ^39–41^; transcription factor EC (*TFEC*), integrin subunit alpha X (*ITGAX*, also known as CD11c), and zinc finger and BTB domain containing 46 (*ZBTB46*) are specific for macrophages and dendritic cells ^42–44^; and follicular DCs are known to robustly express complement C3b/C4b receptor 1 (*CR1*), complement C3d receptor 2 (*CR2*), and C-X-C motif chemokine ligand 13 (*CXCL13*) ^45,46^.

In parallel to the clustering and assignment of the immune cell lineages, we assigned two epithelial clusters based on the airway epithelium lineage hierarchy described by Montoro and colleagues ^47^. Notably, neither the basal/hillock lineage nor club lineage expressed immune cell-related markers and DEGs (**Figure 1C**, **Supplemental Figures 1A, B**). To evaluate the specific cell composition of each epithelial-related lineage cluster, we obtained the module score of a list of characteristic genes of the cell types contained in each lineage (**Figures 1D, E**). Module scores were calculated by normalizing the expression values of each gene to the maximum expression of the combination of genes across all the nuclei. These module scores showed that nuclei identified as reflecting the basal/hillock lineage clearly expressed genes such as insulin like growth factor binding protein 3 (*IGFBP3*), collagen type XVII alpha 1 chain (*COL17A1*), keratin 8 (*KRT8*), keratin 13 (*KRT13*), proteasome 26S subunit, non-ATPase 5 (*PSMD5*), and SRY-box transcription factor 9 (*SOX9*) that have been previously associated with basal, hillock, neuroendocrine, and tuft cells ^47,48^. The club lineage cluster showed higher module scores for unique markers representing club or secretory (secretoglobin family 1A member 1, *SCGB1A1*), ciliated (cadherin related family member 3, *CDHR3*, and forkhead box J1, *FOXJ1*), goblet (mucin 5AC/5B, *MUC5AC/5B*), and microfold (TNF receptor superfamily member 11a, *TNFRSF11A*) cells ^47,49^, than any other cluster.

Due to our interest in the role of epithelial cells in the initiation and development of mucosal immunity in the airway, we focused our analysis on the two epithelial cell lineages present in human adenoids.

### Characterization of basal/hillock lineage cell populations

Having first identified the basal/hillock lineage cluster, we next turned our attention to sub-clustering this lineage into individual cell types using the same bioinformatic pipeline as above. We performed UMAP reduction (dimensionality = 11, resolution = 1) to identify basal cell progenitors, neuroendocrine, tuft, and hillock cells using specific cell marker genes for these cell types (**Figure 2A**, **Supplemental Figures 1C, D**). Genes that have been associated with basal cells such as insulin like growth factor binding protein 3 (*IGFBP3*), delta like non-canonical Notch ligand 2 (*DLK2*) and laminin subunit beta 3 (*LAMB3*) ^48,50,51^ were expressed in basal cluster but also in neuroendocrine and tuft cells. Neuroendocrine cells expressed proteasome 26S subunit, non-ATPase 5 (*PSMD5*), the neurotransmitter-associated gene nebulin (*NEB*), and the peroxisomal biogenesis factor 5 like (*PEX5L*) ^47,50^. Annexin A4 (*ANXA4*), Spi-B transcription factor (*SPIB*), and SRY-box transcription factor 9 (*SOX9*) were significantly expressed in tuft cells ^47,50^. The ‘hillock cells’ that were recently identified in both mouse and human airways were also present in our human adenoid analysis and showed expression of keratin 4 (*KRT4*), sciellin (*SCEL*), and small proline rich protein 1B (*SPRR1B*) ^47,50,52^. Finally, we were unable to assign one cluster to any previously described cell type although it expressed basal-related gene markers (**Figure 2B**, **Supplemental Figure 1D**), and thus for the purpose of downstream analysis we named this cluster “unknown cell type” (UCT).

**Figure 2.**
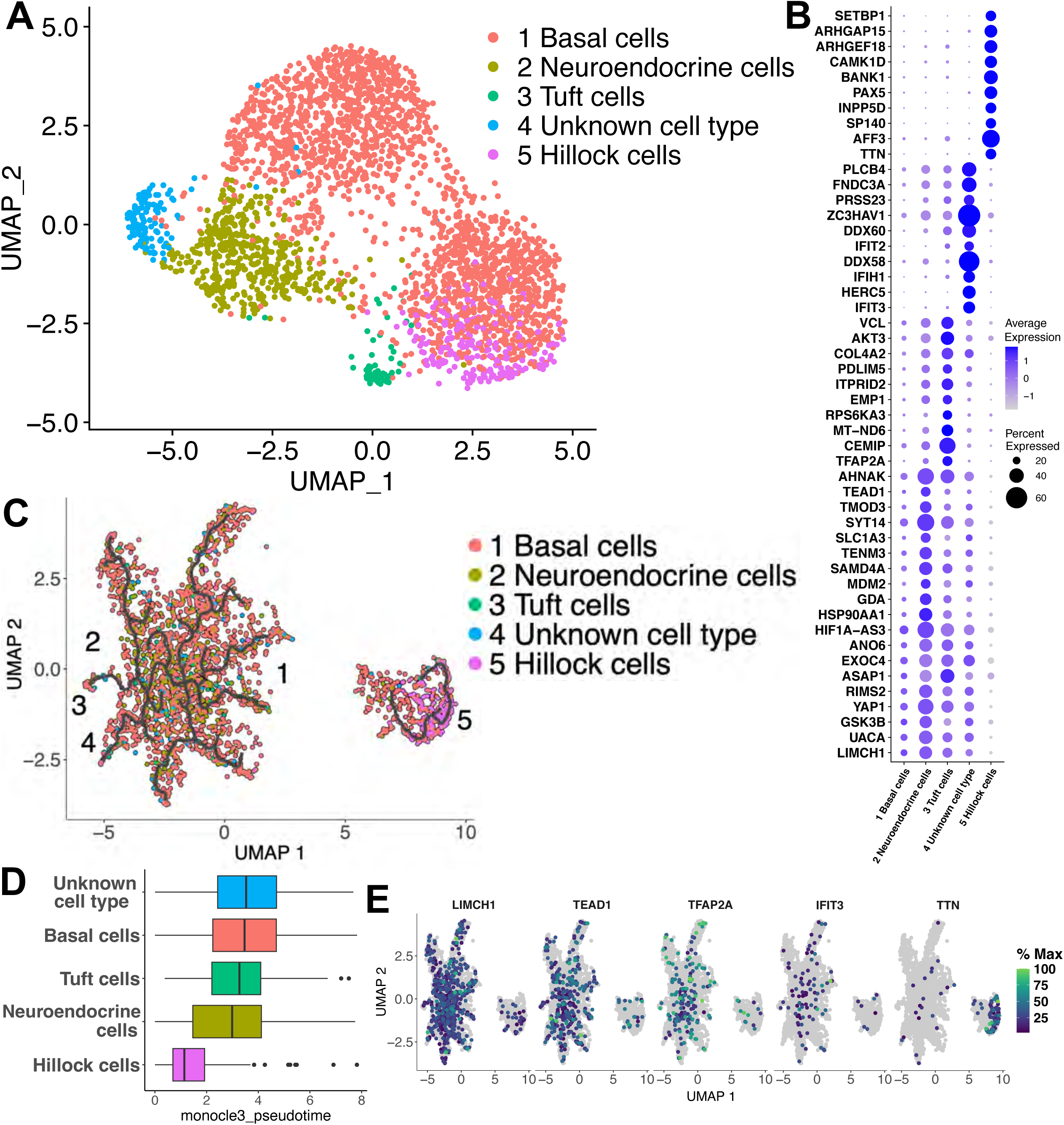
Characterization of basal/hillock lineage cell populations. (A) UMAP plot of the 2,796 nuclei from subset basal/hillock lineage cluster of the human adenoids (dim. = 11, res. = 1). (B) Dot plot of the average expression of top DEGs for each nucleus cluster in this lineage. (C) Trajectory analysis with Monocle 3 of the clusters identified in the basal/hillock lineage. The black line in each group of nuclei represents the different trajectories identified by the pipeline. (D) Nucleus clusters organized by pseudotime setting as root nodes the ones specified in the Sup. Fig2A. One separated root node was designated for each trajectory group. The black bar inside each box represents the median value of the data. (E) Percentage of maximum expression of a representative DEG of each cluster through the differentiation pathway.

After labeling the different nucleus populations by using previously described cell marker genes, we analyzed the average expression of the top DEGs for each of the 5 clusters (**Figure 2B**). Notably, as we observed with the basal-associated gene markers (**Supplemental Figures 1C, D**), DEGs found in the basal cell sub-cluster were also expressed by neuroendocrine and tuft cells (**Figure 2B**). This observation supported that neuroendocrine and tuft cells differentiate from basal progenitor cells^47^. The cell cluster that we named UCT not only expressed basal cell DEGs but also had a unique and unexpected transcriptional signature highlighted by robust expression of several interferon-stimulated genes (ISGs) (**Figure 2B**). Finally, the ‘hillock cells’ did not express DEGs characteristic of any basal cell-related cluster and had a distinct transcriptional signature (**Figure 2B**).

To determine if the basal-related clusters and the hillock cluster have a common cell progenitor, we applied the Monocle 3 analytical pipeline ^53,54^ to plot the single-nucleus trajectories of all identified clusters. This analysis generated a UMAP plot with two clearly distinct nucleus clusters (**Figure 2C**). While all the basal-cluster derived cells were grouped together, nuclei representing the hillock cluster were divergent. The Monocle 3 bioinformatic analysis identified two different trajectory pathways, one for each group of cells. After manually selecting the root nodes for each trajectory, we determined the distance of each nucleus to the differentiation pathway origin through pseudotime analysis (**Supplemental Figure 3A**). Notably, a higher pseudotime value for any given nucleus predicts that the cell originated later in the differentiation pathway. After organizing nucleus clusters by pseudotime, we observed that hillock cells originated significantly earlier than the other cell clusters (**Figure 2D**). In contrast, the basal-related clusters had similar median pseudotime values, reflecting their shared differentiation from basal cells.

Finally, we used Monocle 3 to analyze the expression variability of a representative DEG for each cell cluster (**Figure 2B**) along the trajectory (**Figure 2E**). We decided to use DEGs instead of specific cell marker genes for this analysis because of their absence from the UCT cluster and the changing list of markers for the recently identified hillock cells. While all the basal-related cluster DEGs were mostly expressed across the cell group that contained basal-cluster derived cells, the hillock DEG titin (*TTN*) showed significant expression only in the hillock cell cluster, highlighting once again their different origin. Similarly to the trajectory UMAP plot (**Figure 2C**), the basal-related cluster DEGs showed expression patterns that mostly overlapped with each other, and did not show a unique differentiation pathway for each corresponding cell cluster (**Figure 2E**).

Taken together, our analysis of the basal cell lineage allowed us to identify and define the transcriptional landscape of basal cells in the human adenoid, their progeny, as well as the recently described hillock cells. Moreover, we unexpectedly identified a cluster of cells with a unique transcriptional signature that we named “unknown cell type”.

### The “unknown cell type” is a basal cell progeny with a unique interferon-related gene signature

During our subcluster analysis of the basal/hillock lineage, we identified a cluster of cells derived from basal cells that did not share either cell marker genes or DEGs with previously described basal-related cell types (**Figure 2B**, **Supplemental Figure 1D**). When we analyzed the individual expression levels in transcripts per million (TPM) of the top DEGs representing this UCT population and compared them to the other basal/hillock lineage cells, the differences were clearly visible (**Supplemental Figure 4A**). The unique expression profile of the UCT cells was even more apparent when we assessed UCT-specific DEG expression across all the nucleus populations identified in the human adenoid (**Figure 3A**). In this analysis, we included not only the basal/hillock lineage but also the cell types from the club lineage analysis that will be described below. From these analyses, a gene signature including the transcripts for interferon induced proteins with tetratricopeptide repeats 2 and 3 (*IFIT2* and *IFIT3*), the HECT and RLD domain containing E3 ubiquitin protein ligase 5 (*HERC5)*, the RNA sensor RIG-I (*DDX58*), and the DExD/H-box helicase 60 (*DDX60*) were uniquely characteristic of the UCT population.

**Figure 3.**
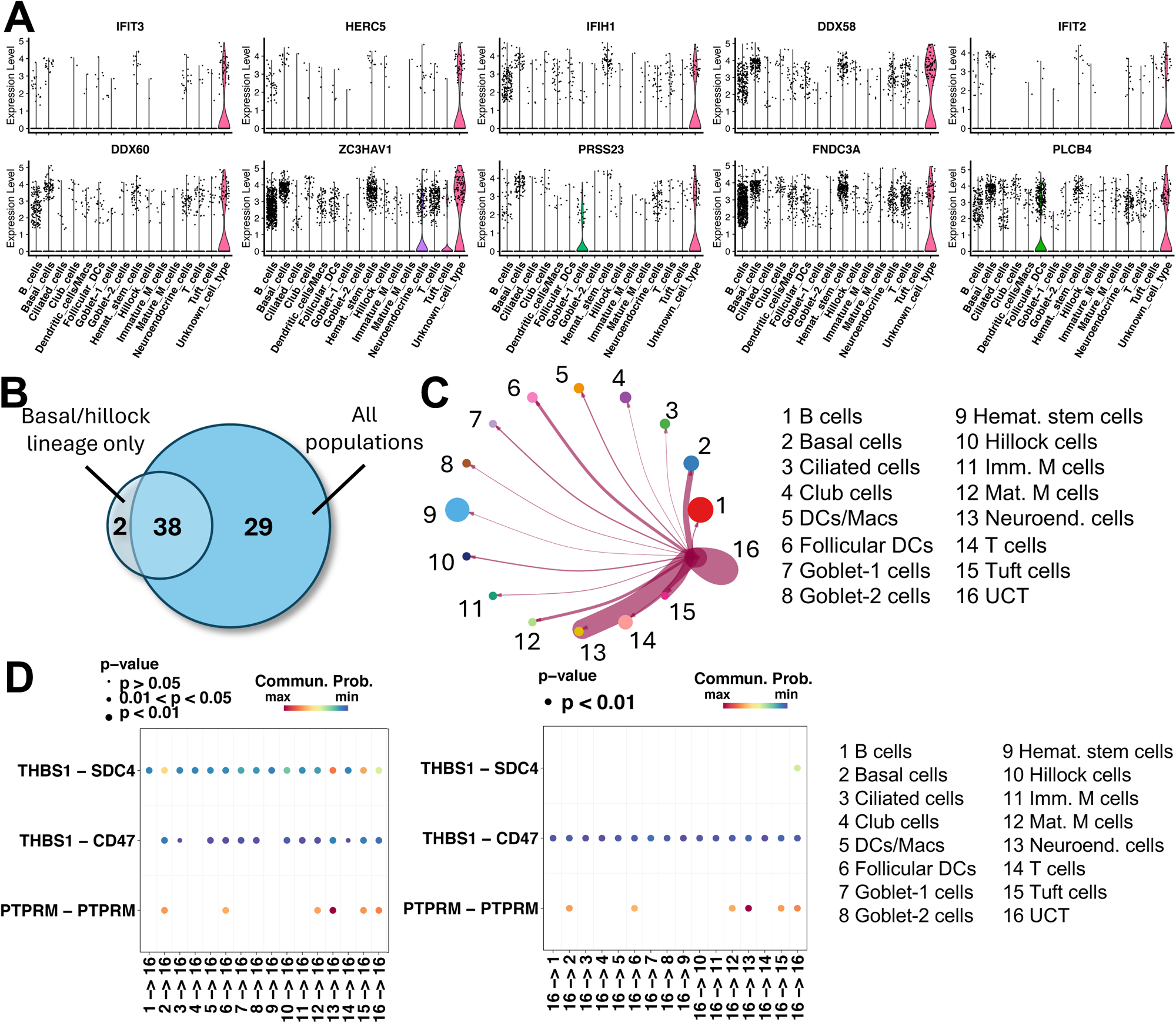
The “unknown cell type” is a basal cell progeny with a unique interferon-related gene signature. (A) Individual expression levels in transcripts per million (TPM) of the top 10 UCT DEGs in all the nucleus populations identified in the human adenoid. (B) Venn diagram showing overlap between UCT DEGs identified only using basal/hillock lineage populations or all the human adenoid populations. (C) Cell-cell communications of UCT cells with all the other cell types present in the human adenoid. The size of the colored circle of each cluster is proportional to the number of nuclei in each. The thickness of each connecting line is proportional to the predicted likeliness of the interaction between a certain pair of cell clusters. (D) Communication probability of the predicted most significant ligand-receptor interactions of all cell types in the human adenoid with UCT cells (left) and vice versa (right). The p-value, communication probability and number corresponding to each cluster are indicated in the legend of the bubble plots.

Next, we attempted to identify markers to discriminate UCT cells not only from the basal lineage populations but also from all cell populations in the human adenoid. The new list of DEGs for UCT mostly overlapped with the one obtained from the basal/hillock lineage analysis (**Figure 3B**) and the top DEGs were almost identical (**Supplemental Figure 4B**).

Though we were clearly struck by the abundant expression of interferon-induced genes, we next used an unbiased pathway analysis to further categorize the unique gene signature of these cells. Using the Database for Annotation, Visualization, and Integrated Discovery (DAVID) tool ^55,56^, we identified biological processes in which the DEGs of the UCT population were enriched and sorted those processes into three groups (**Table 1**). The first group included processes involved with cell signaling like cell morphogenesis, regulation of cellular process, response to stimulus, and both cell-cell and intracellular signaling. The second group included biological processes involved in cellular responses to infection. The third group of biological processes were classified for their roles in migration and differentiation. It should be noted that, though these adenoids were collected from donors undergoing elective adenoidectomy for sleep apnea, we cannot exclude the possibility that the donors were experiencing a coincidental viral or bacterial infection at the time of surgery, thus eliciting an interferon response from an otherwise quiescent cell.

**Table 1:** Identification of the biological processes important for the novel interferon-related “unknown cell type” identified in the human NALT. Only processes with a p-value < .05 are considered and grouped in the different categories.

|  |  | **Gene annotated count** | **p-Value** |
| --- | --- | --- | --- |
| **Signaling** | Cell morphogenesis | 19 | 1.2E-09 |
|  | Regulation of cellular process | 53 | 5.1E-06 |
|  | Response to stimulus | 45 | 8E-05 |
|  | Cell-cell signaling | 15 | 3.2E-04 |
|  | Intracellular signal transduction | 21 | 1.4E-04 |
| **Infection response** | Response to stress | 32 | 2.2E-07 |
|  | Regulation of cell-substrate adhesion | 7 | 6.6E-05 |
|  | Response to virus | 7 | 1.5E-03 |
|  | Immune system process | 20 | 1.8E-03 |
| **Migration and differentiation** | Cell differentiation | 35 | 1.7E-08 |
|  | Cell adhesion | 19 | 6.8E-07 |
|  | Cell motility | 18 | 2.5E-05 |
|  | Locomotion | 19 | 3.2E-05 |
|  | Regulation of cell migration | 13 | 7.8E-05 |
|  | Epithelial cell development | 5 | 7.7E-03 |

To further understand unique characteristics of the UCT cell cluster, we used the CellChat pipeline ^57^ to predict cell-cell communication based on the unique expression of genes among the cells within the tissue and to identify potential interactions of the UCT with other cell types. We first used CellChat to predict the interactions of UCT cluster with any other cluster individually (**Figure 3C**). Here, the thickness of each line is proportional to the predicted likelihood of an interaction between pairs of cells ^57^.

The strongest interactions predicted for UCT cells were with themselves, neuroendocrine, tuft, basal, follicular DCs, and mature M cells, with the greatest predicted interaction with neuroendocrine cells (**Figure 3C**). CellChat can also predict receptor-ligand interactions based on gene expression data ^57^. Thus, we determined that the protein tyrosine phosphatase receptor type M (PTPRM) and thrombospondin (THBS) pathways have the strongest communication probabilities for mediating interactions of UCT cells with other cell types and vice versa (**Supplemental Figure 4C**). PTPRM is a member of the type IIb subfamily of receptor protein tyrosine phosphatases that has important roles in development ^58^, cell-cell adhesion ^59^, motility ^60^, and oncogenesis ^61^. Also, the head-to-tail homodimers have been predicted to facilitate other ligand-receptor interactions between two cell membranes ^59^. CellChat predicted that the strongest interaction of UCT cells was the self-interaction of PTPRM between UCT cells, along with PTPRM interactions between UCT cells and other cell types (**Figure 3D**). Interaction between thrombospondin 1 (THBS1) and syndecan 4 (SDC4) was predicted to mediate the interaction of basal, neuroendocrine, and tuft cells with UCT cells (**Supplemental Figure 4D**). Although UCT cells demonstrated low levels of *THBS1* expression, the reciprocal interaction of THBS1 and SDC4 may explain the high interaction probability of this pair within the UCT cells. Finally, despite UCT cells displaying high expression of genes related to interferon responses, we were unable to identify specific pathways through which cell surface interferon receptors could impact interactions of UCT cells with either epithelial or hematopoietic lineage cells within the adenoid.

Taken together, our analysis revealed the presence of a unique cell type in the basal cell lineage that demonstrated abundant expression of interferon related genes, a relevant role in cellular responses to infection, and unique receptor ligand interactions with other adenoid cell populations.

### Characterization of the club lineage cell populations

As with the basal/hillock lineage, we re-analyzed the club lineage cells to identify individual cell populations. The UMAP plot (dimensionality = 11, resolution = 0.9) for club lineage cells allowed us to label unique nucleus populations by using previously published data for markers of specific club cell populations (**Figure 4A**, **Supplemental Figures 1E, F**). We first identified a cluster of club cells, also named secretory or Clara cells ^25^, representing the precursor cell for all cells in this lineage (**Figure 4A**). Despite not demonstrating a significant expression of club marker genes secretoglobin family 1A member 1 (*SCGB1A1*) or nuclear factor I A (*NFIA*) ^47,50^ (**Supplemental Figure 1F**), that club DEGs were expressed by all other clusters in this lineage and their distribution in the trajectory analysis that we detail below (**Figures 4B, C**), guided us to label this cluster as club cells. Of note, very low expression of *SCGB1A1* has also been observed previously in nasal secretory cells ^50^. We next identified a cluster of ciliated cells by expression of genes related to cilia formation and function ^62–64^ including cadherin related family member 3 (*CDHR3*), the forkhead box J1 (*FOXJ1*), and sentan, cilia apical structure protein (*SNTN*) ^47,50^. This cluster also included other DEGs (**Figure 4B**), beside the aforementioned *CDHR3* gene, the RP1 axonemal microtubule associated (*RP1*), cilia and flagella associated protein 54 (*CFAP54*), and dynein axonemal heavy chain 12 (*DNAH12*) that were annotated in the ciliated cluster of a recent snRNA-seq analysis of mouse lungs ^18^. Also, we classified two variants of goblet cells: a goblet-1 cluster that expressed mucin 5AC/5B (*MUC5AC/5B)* ^47^, and a goblet-2 cluster that expressed mucin 4 (*MUC4*), interleukin 19 (*IL19*), and colony stimulating factor 3 (*CSF3*) ^47,65^. Beside these mucin family members, DEGs for both goblet cell populations included other genes including BPI fold containing family member 1 (*BPIFB1*) ^66^, polymeric immunoglobulin receptor (*PIGR*) ^67^, serine protease 23 (*PRSS23*) ^68^, and lipocalin 2 (*LCN2*) ^69^, that have also been associated with mucus production and secretion in human and murine airways ^47,52^ (**Figure 4B**). Notably, both goblet cell clusters shared many of their DEGs though with varying expression levels (**Figure 4B**), highlighting their close ontology within the lineage. Finally, we identified two populations of M cells that we classified as immature and mature cells based on the expression of M cell markers consistent with immature (*RELA/B* NF-KB subunits, SRY-box transcription factor 8, *SOX8*) or mature (*SPIB*, TNF receptor superfamily member 11a, *TNFRSF11A*, and TNF-a induced protein 2, *TNFAIP2*) ^13,49,70^ M cells, and their position in trajectory analysis (**Figure 4C**). Immature M cells were characterized by DEGs that demonstrated lower average expression levels compared to mature M cells (**Figure 4B**). Notably, the existence of a population of immature or transitional M cells during their differentiation and maturation has been described previously in mouse GI tract and intestinal organoids ^71,72^. Mature M cells had a distinct gene signature that we discuss in greater detail below.

**Figure 4.**
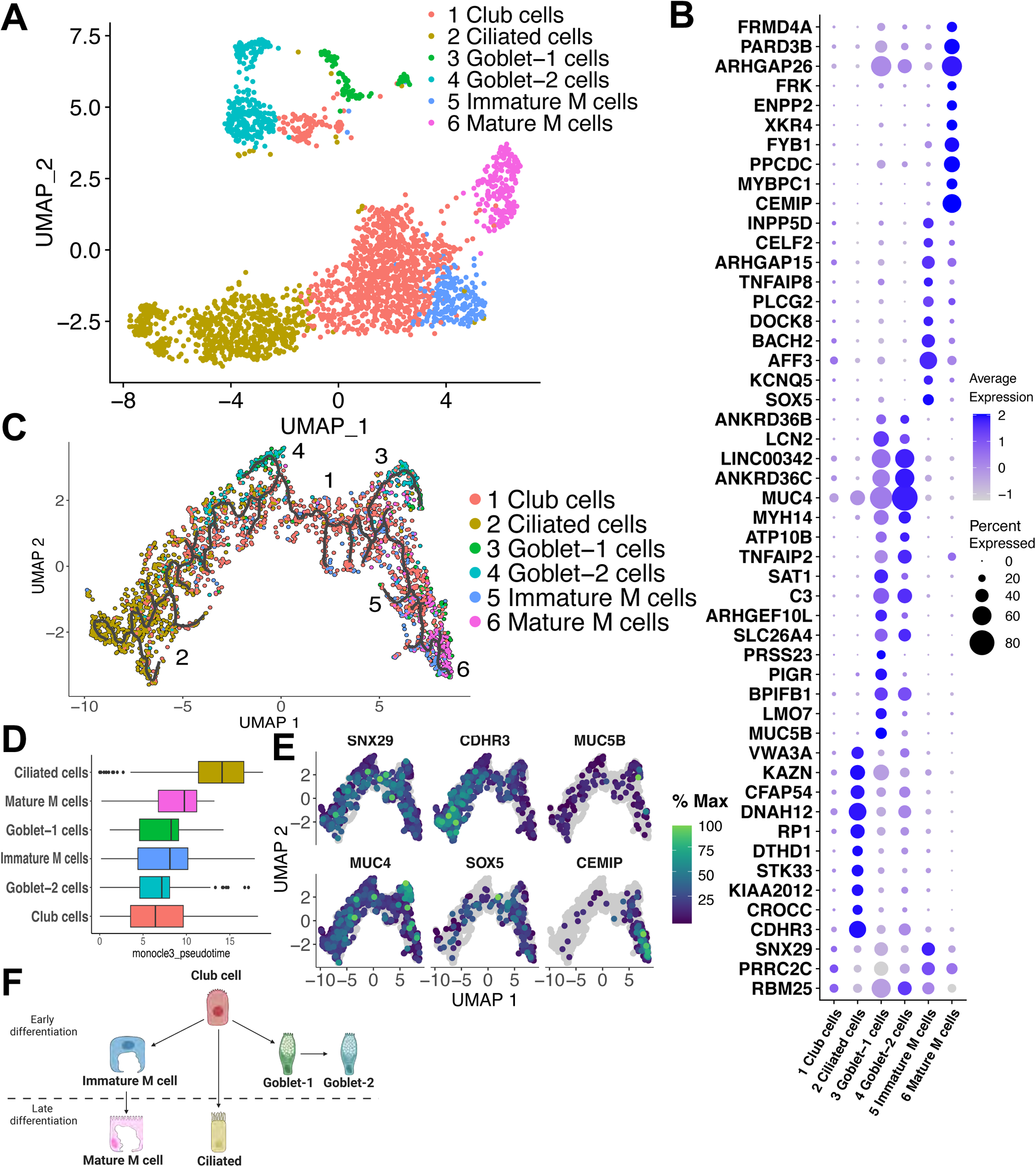
Characterization of the club lineage cell populations. (A) UMAP plot of the 2,469 nuclei from subset club lineage cluster of the human adenoids (dim. = 11, res. = 0.9). (B) Dot plot of the average expression of top DEGs for each nucleus cluster in this lineage. (C) Trajectory analysis with Monocle 3 of the clusters identified in this lineage. The black line represents the trajectory identified by the pipeline. (D) Nucleus clusters organized by pseudotime setting as root node the one specified in the Sup. Fig2B. The black bar inside each box represents the median value of the data. (E) Percentage of maximum expression of a representative DEG of each cluster through the differentiation pathway. (F) Club cell lineage hierarchy obtained from the analysis of the DEG and trajectory results.

To verify that our cell cluster labeling fit with a previously described club cell hierarchy ^47^, we applied single-nucleus trajectory analysis to the club lineage clusters using the Monocle 3 pipeline ^53,54^. Trajectory analysis generated a UMAP plot containing a single trajectory that connected all the cell types through their differentiation (**Figure 4C**). Club cells were predominantly centrally located in the trajectory, but also were distributed in both vectors of the differentiation pathway. Ciliated cells were enriched to the left side of the trajectory while immature and mature M cells were enriched on the right side. The two populations of goblet cells overlapped, were centrally located and symmetrical on the differentiation pathway. After manually selecting the root node and determining pseudotime values for each cell population (**Supplemental Figure 3B**), we then organized the cell clusters by pseudotime (**Figure 4D**). With this analysis we determined that club cells, despite being dispersed throughout the trajectory (**Figure 4C**), are the earliest appearing cells in the lineage. With similar medium pseudotime values, the next cells in the differentiation pathway were the two goblet cell populations and immature M cells. Finally, mature M cells and ciliated cells had larger median pseudotime values indicative of a later appearance, although this could also be due to a high turnover ^73^.

To further highlight the unique differentiation pathways of the various cell types, we chose a representative DEG for each cell type to demonstrate either its pleiomorphic or distinct expression pattern as we did for the basal/hillock lineage (**Figure 4E**). While the club cell lineage representative gene *SNX29* was expressed in cells dispersed across the differentiation trajectory, its maximum expression was in central cells (**Figure 4E**). *CDHR3*, a marker of ciliated cells, demonstrated increasing expression in cells distributed on the left side of the trajectory consistent with the position of ciliated cells (**Figures 4C, E**). *MUC5B* and *MUC4*, gene markers and DEGs for goblet-1 and goblet-2 cells respectively, had overlapping expression distributed in both arms of the trajectory (**Figure 4E**), highlighting the secretory functions of these cell clusters and also of other clusters of this lineage ^50^. The expression pattern of *SOX5*, previously reported as part of a complex regulatory network with other Sox proteins including the early M cell marker *SOX8* ^74^, predominantly overlapped in the region of the trajectory containing immature M cells (**Figures 4C, E**). Finally, *CEMIP*, a gene that in our analysis was differentially expressed in mature M cells within the club cell lineage demonstrated maximum expression at the right terminus of the differentiation pathway where mature M cells were found (**Figures 4C, E**).

We propose a lineage model where club cell progenitors differentiate first into immature M cells and goblet cells. Whereas immature M cells then differentiate into mature M cells, ciliated cells differentiate directly from club cells over a longer time course (**Figure 4F**). The high degree of overlap between both goblet cell populations in their DEG expression and trajectory analysis suggest that both cell types are not directly differentiated from club cell progenitors. Instead, we proposed that goblet-1 cells differentiate from club cells as a transition to the goblet-2 cell population.

Taken together, our analysis of the club lineage cell types allowed us to not only identify two populations of M cells in the human adenoid, but also determine their differentiation path.

### Characterization of human adenoid M cells

After identifying and labeling two populations of nuclei as M cells (**Figure 4A**, **Supplemental Figures 1E, F**) we next determined the expression of previously described M cell markers in all adenoid cell populations. Because no data are available for human adenoid M cells, we collated a list of potential markers from human GI tract cells or organoid M cells ^13,72,75–78^, or from recent studies of mouse lung M cells ^17,18^ (**Figure 5A**). We began our analysis by assessing previously established genes including the transcription factors *SOX8* and *SPIB*, cell surface protein *GP2*, the C-C motif chemokine ligand 20 (*CCL20*), and the TNF family signaling pathway components *TNFRSF11A*, *TNFRSF11B*, and *TNFAIP2*. The transcription factor SOX8 has been described as essential for the differentiation of M cells and antigen-specific IgA response in the GI tract ^71^. Although the percent expression of *SOX8* was low within the cells of the adenoid, mature M cells had the highest average expression amongst the cells. *SPIB*, a transcription factor involved in M cell differentiation both in the GI tract ^13,72,75,79–81^, and airway MALT ^17,18^, was highly expressed in the mature human adenoid M cells. As expected, B cells, DCs/macrophages, follicular DCs ^82,83^, immature M cells, and tuft cells ^47^ also expressed *SPIB*, albeit with fewer cells and reduced expression. The chemokine CCL20 was previously described as specific for M cells in Peyer’s patches ^78^, but also expressed by goblet cells at mucosal surfaces ^84^. In the adenoid, while less than 10% of mature M cells expressed *CCL20*, they did have the highest expression levels. Another cell surface protein traditionally considered a specific M cell marker in both mouse and human tissues is glycoprotein 2 (*GP2*) ^85^, although GP2 is not expressed in *in vitro* M cell models ^15^, and some authors have reported that GP2 may not be specific for nasal and ocular M cells in mice ^86^. In our dataset, very few cells expressed *GP2*, suggesting it may not be abundant in human adenoid MALT tissues. Similarly, the interaction between TNF receptor superfamily member 11a (TNFRSF11A, also known as RANK) and its ligand TNF superfamily member 11 (TNFSF11 or RANKL) is necessary for differentiation but insufficient for full maturation of M cells in the GI tract and mouse NALT ^75,86–88^. Furthermore, soluble TNFSF11 (RANKL) has been used to induce M cell differentiation *in vitro* ^72,86^. Likewise, mature M cells produce TNF receptor superfamily member 11b (TNFRSF11B, also named OPG) to inhibit the differentiation of epithelial cells into M cells ^77^. In our analysis, the percentage of either immature or mature M cells that expressed either *TNFRSF11A* (*RANK*) or *TNFRSF11B* (*OPG*) was very low (**Figure 5A**). Lastly, *TNFAIP2*, whose expression is induced by SPIB ^13,71,75^ and functions in the exocytosis process in M cells ^76,86,89,90^, was significantly expressed in both goblet cells and mature M cells. Overall, while we confirmed expression of *SPIB, TNFRSF11A* and *TNFAIP2* in mature adenoid M cells, transcripts for other previously described genes were absent.

**Figure 5.**
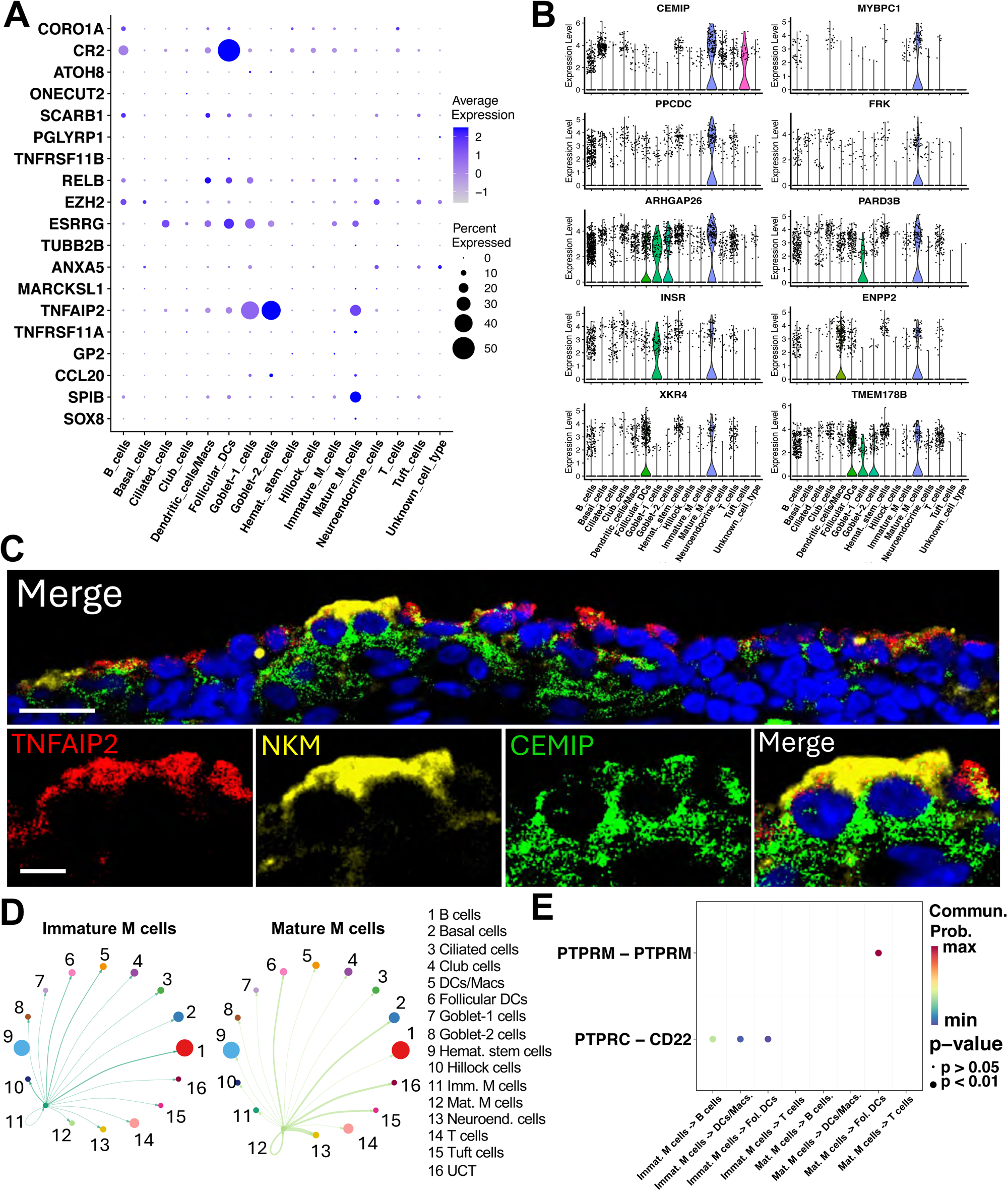
Characterization of human adenoid M cells. (A) Dot plot of the average expression of previously described M cell markers in all nucleus populations identified in the human adenoid. (B) Individual expression of the top 10 DEGs for mature M cells in all nucleus populations identified in the human adenoid. (C) Human adenoid sections were stained with anti-TNFAIP2 antibody, NKM 16-2-4, and anti-CEMIP antibody, and analyzed by confocal microscopy. Scale bar, top 15 μm, bottom, 5 μm. (D) Cell-cell communications of immature (left) and mature (right) M cells with all the other cell types present in the human adenoid. The size of the colored circle of each cluster is proportional to the number of nuclei in each. The thickness of each connecting line is proportional to the predicted likeliness of the predicted most significant interaction between a certain pair of cell clusters. (E) Communication probability of the ligand­-receptor interactions of all immune cell types in the human adenoid with immature (left) and mature (right) M cells. The p-value, and communication probability are indicated in the legend.

Although we initially assembled a list of DEGs for mature M cells during our analysis of club lineage cells (**Figure 4B**) that clearly differentiated them from the other cells in the lineage (**Supplemental Figure 5A**), we next generated a parallel panel of DEGs for mature M cells using all the nucleus populations we identified in adenoid inclusive of immune and basal cell lineages. When we compared the two gene sets, we found that most genes unique to mature M cells were identified in both analyses (**Supplemental Figure 5B**). However, 13 genes were unique to the analysis performed with club lineage cells while 5 were uniquely identified in comparison to all the cells (**Supplemental Figures 5B, C**). Of the top ten DEGs from the all-cell analysis, only two differed from the prior club lineage analysis (**Figure 5B**). We identified several genes with potential functional relevance to mucosal immunity. The cell migration inducing hyaluronidase 1 (CEMIP) is involved in degradation of the long chain glycosaminoglycan hyaluronan ^91^ and plays a role in regulating skin immunity to *Staphylococcus aureus* infection ^92^. The myosin binding protein C1 (MYBPC1) is a cytosolic protein that interacts with myosin and actin filaments ^93^, and thus may be involved in M cell transcytotic activity. The phosphopantothenoylcysteine decarboxylase (PPCDC) protein has decarboxylase activity and is involved in the production of coenzyme A, an important cofactor in intracellular reactions and critical for lipid biosynthesis for Mtb during infection ^94^. The rho GTPase activating protein 26 (ARHGAP26) has roles in endocytosis, cell spreading, and muscle development ^95^, though a role in mucosal immunity has not been described. The insulin receptor (INSR) is present in multiple cell types and its role in enhancing immunity during inflammation and infection may contribute to the development of mucosal immune response in human NALT ^96^, though a function in M cell biology is not known. The ectonucleotide pyrophosphatase/phosphodiesterase-2 (ENPP2, also known as autotaxin) is involved in the production of lysophosphatidic acid (LPA) that through its interaction with specific G protein-couple receptors (GPCRs) is able to promote cell responses such as migration, proliferation, and survival ^97,98^. Finally, the transmembrane protein 178B (TMEM178B) has been associated with inflammation progression in patients with severe asthma ^99^.

We next used pathway analysis to broadly determine which biologic pathways were enriched in adenoid mature M cells (**Table 2**). Pathways involved in cell adhesion were enriched, consistent with the interconnection of M cells with other MALT cells. Another broad set of pathways were indicative of cell signaling both within M cells but also to other adenoid cells. Moreover, many signaling pathways were predicted to impact or regulate localization of other cells. Finally, the most enriched of all biologic pathways were those involved in trafficking, such as those related to transport, location, or endocytosis, which likely reflects the transcytotic activity of M cells ^15^.

**Table 2:**
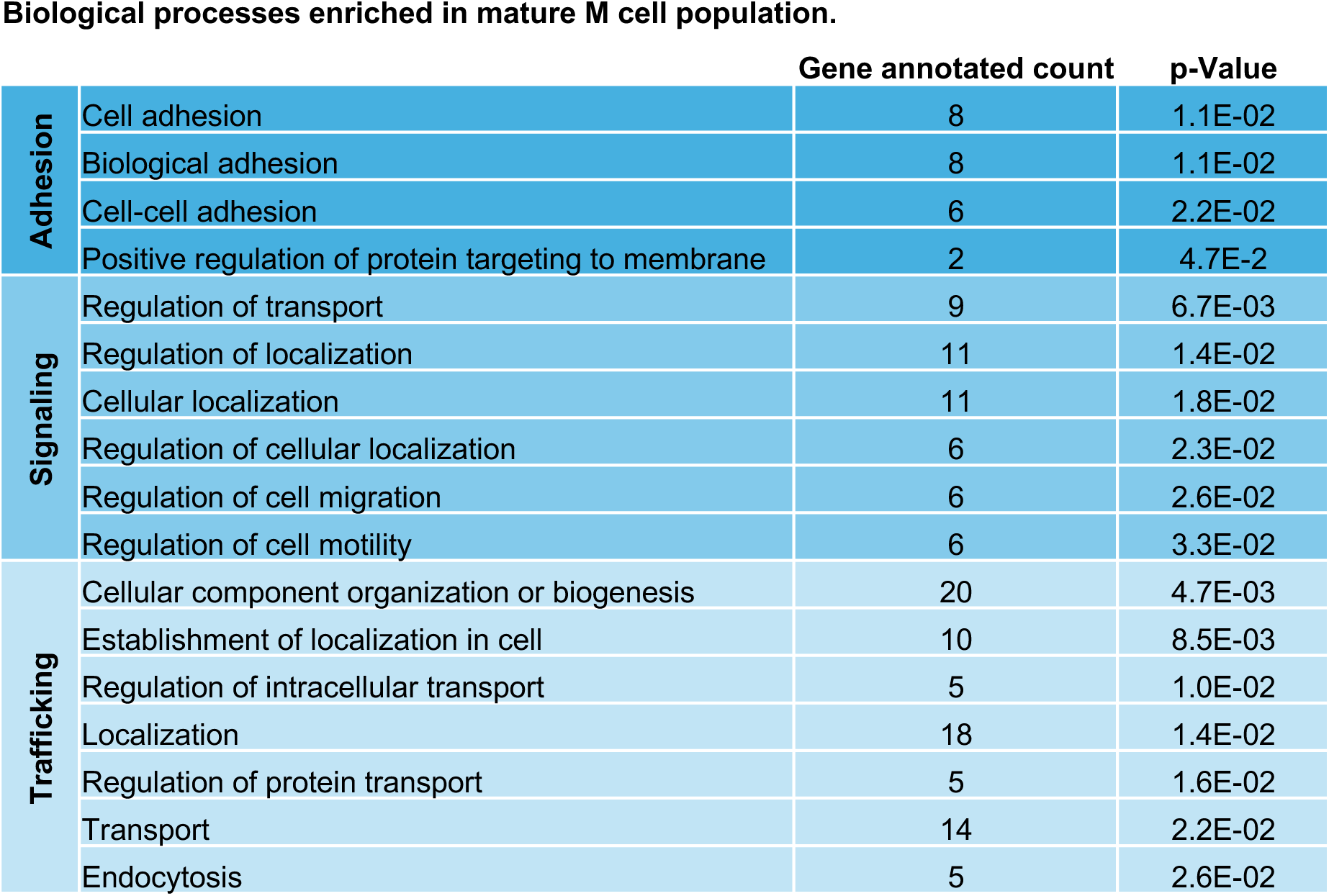
Pathway enrichment analysis for the human adenoid mature M cells. Only processes with a p-value < .05 are considered and grouped in the different categories.

|  |  | Gene annotated count | p-Value |
| --- | --- | --- | --- |
| Adhesion | Cell adhesion | 8 | 1.1E-02 |
|  | Biological adhesion | 8 | 1.1E-02 |
|  | Cell-cell adhesion | 6 | 2.2E-02 |
|  | Positive regulation of protein targeting to membrane | 2 | 4.7E-2 |
| Signaling | Regulation of transport | 9 | 6.7E-03 |
|  | Regulation of localization | 11 | 1.4E-02 |
|  | Cellular localization | 11 | 1.8E-02 |
|  | Regulation of cellular localization | 6 | 2.3E-02 |
|  | Regulation of cell migration | 6 | 2.6E-02 |
|  | Regulation of cell motility | 6 | 3.3E-02 |
| Trafficking | Cellular component organization or biogenesis | 20 | 4.7E-03 |
|  | Establishment of localization in cell | 10 | 8.5E-03 |
|  | Regulation of intracellular transport | 5 | 1.0E-02 |
|  | Localization | 18 | 1.4E-02 |
|  | Regulation of protein transport | 5 | 1.6E-02 |
|  | Transport | 14 | 2.2E-02 |
|  | Endocytosis | 5 | 2.6E-02 |

To conclude the analysis of the mature M cell DEGs, we studied the colocalization of some classical M cell markers and the top DEG on the list, CEMIP, by immunofluorescence microscopy on human adenoid tissue sections (**Figure 5C**). We used staining with the classical markers TNFAIP2 because *TNFAIP2* was expressed in the human mature M cell population (**Figure 5A**, **Supplemental Figure 1F**), and NKM 16-2-4 (a monoclonal antibody specific for a1,2-fucose ^3^) because NKM 16-2-4 has been reported to be a specific marker for M cells ^3^. Confocal microscopy demonstrated colocalization of CEMIP with cells positive for NKM 16-2-4 and TNFAIP2 in the human adenoid (**Figure 5C**).

We next analyzed cell-cell interactions of M cells with all other adenoid cells. To determine how the transition from an immature to a mature M cell impacts cell-cell interactions, we studied each M cell cluster independently (**Figure 5D**). Immature M cells established their strongest interactions with B cells, DCs and macrophages, follicular DCs, and hillock cells. In contrast, mature M cells had their strongest interactions with basal cells, follicular DCs, neuroendocrine cells, tuft cells, UCT cells, and with other mature M cells. As immune cells are critical for M cell development and function ^15,49^, we endeavored to identify ligand-receptor interactions that may mediate M cell biology. In the analysis we included the interactions of immature and mature M cells with B cells, DCs and macrophages, follicular DCs, and T cells and discarded low probability interactions (**Figure 5E**). Surprisingly, we did not identify the interaction between *TNFRSF11A* (*RANK*) and *TNFSF11* (*RANKL*) among the most likely interactions predicted by the CellChat pipeline for adenoid M cells likely due to the very low levels of expression of *TNFRSF11A* by adenoid M cells (**Figure 5A**). The main predicted interaction of immature M cells was through the protein tyrosine phosphatase receptor type C (PTPRC) with CD22, also called SIGLEC2, on both B cells and hillock cells. In contrast, the PTPRC-CD22 interaction was not predicted between immature M cells and T cells due to the low expression of both receptors by T cells (**Supplemental Figure 5D**). Very low expression of transcripts for both surface receptors by mature M cells likely explained why this interaction was not identified between mature M cells and immune cell types. CellChat also predicted a homologous interaction of the protein tyrosine phosphatase receptor type M (PTPRM) as driving a strong interaction of mature M cells with follicular DCs. Further analysis of *PTPRM* expression by other nucleus clusters identified that the PTPRM-PTPRM interaction may also be responsible for the strong interaction of mature M cells with basal cells, neuroendocrine cells, tuft cells, UCT cells, and other mature M cells.

Taken together, using a transcriptomics bioinformatics pipeline we characterized two unique M cell clusters -immature and mature-, their prominent molecular and cellular functional pathways, and their primary interactions with other cells of the human adenoid.

### Comparative analysis of human adenoid M cell DEGs

To conclude the analysis of the human adenoid M cell populations, we compared expression of M cell DEGs identified here to prior cell atlas studies of airway and GI tract cells ^17–19,72^. Recently, a cell atlas of the human palatine tonsil, a MALT structure of the Waldeyer’s ring, was described ^19^. M cells have been identified in human tonsil through ultrastructural analyses and immunohistochemistry ^100–103^. Though an M cell population in the human palatine tonsil was not specifically identified in the prior study ^19^, expression of classical M cell markers *SPIB* and *MARCKSL1* was reported in an epithelial cell population named ‘crypt cells’ ^19^. When we interrogated all human tonsillar epithelial cells, we found that the majority of mature M cell DEGs from our analysis in the adenoid were expressed by ‘crypt cells’, although some adenoid M cell DEGs were absent such as *CEMIP*, *XKR4* and *TMEM178B* (**Figure 6A**). Notably, the percentage of crypt cells that expressed mature M cell DEGs was very low (typically below 20%), while other tonsillar populations more robustly expressed some M cell markers. In the case of the immature M cell DEGs, we found that only *DOCK8*, *PLGC2*, and *TNFAIP8* were significantly expressed in the dataset by almost all the human epithelial cell populations in the tonsil.

**Figure 6.**
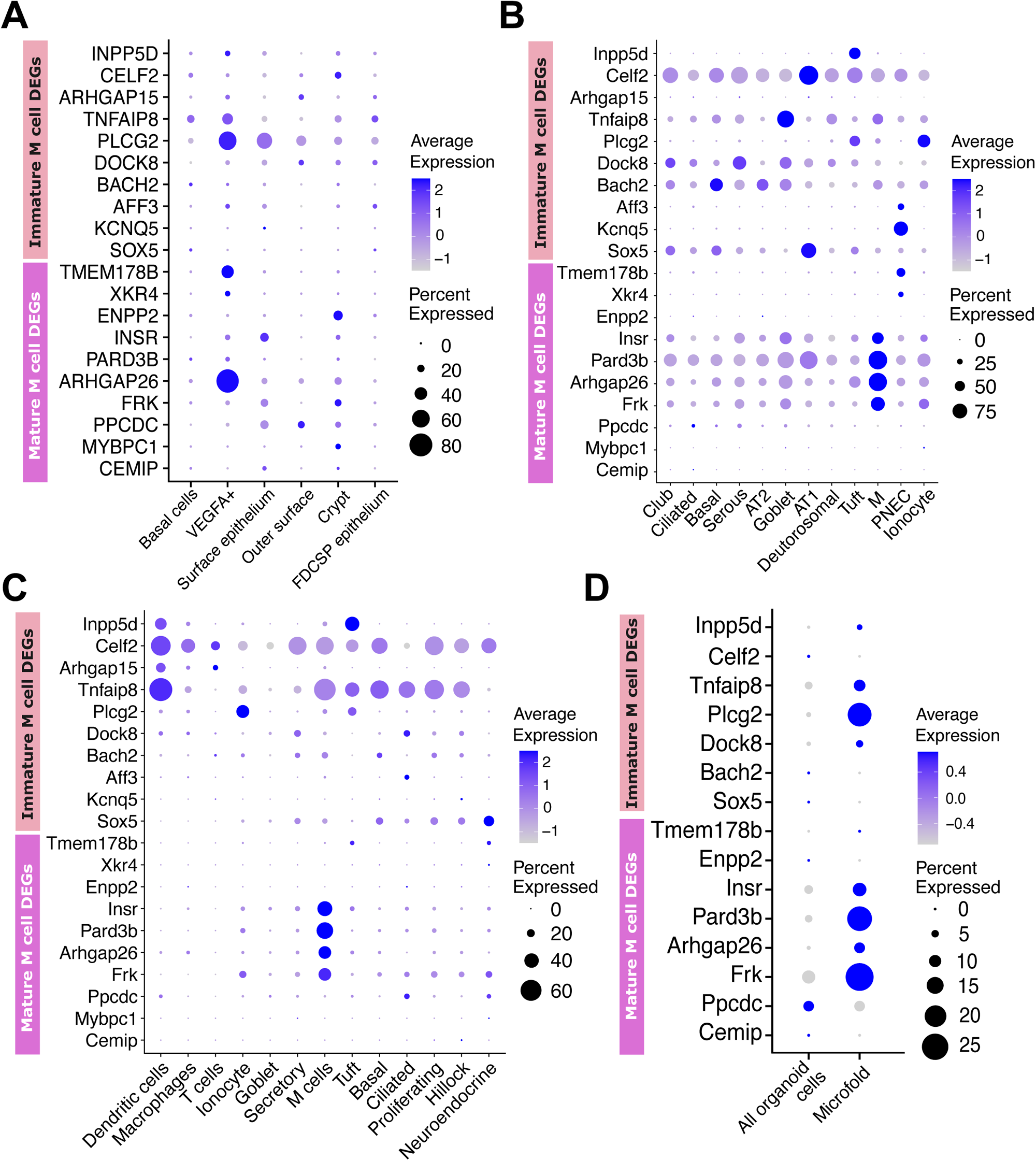
Comparative analysis of human adenoid M cell DEGs. Dot plots of the average expression of newly identified DEGs of mature and immature M cell populations in human epithelial tonsillar cells from Massoni-Badosa et al. 2024 (A), mouse lungs from Barr et al 2023 (B), murine tracheal cells from Surve et al. 2023 (C), and mouse gut organoids treated with RANKL from Haber et al. 2017 (D).

Next, we compared our data to atlases describing M cells in mouse lungs ^18^ (**Figure 6B**), and in mouse tracheal cells ^17^ (**Figure 6C**). We identified a group of mature M cell DEGs -*FRK*, *ARHGAP26*, *PARD3B*, and *INSR*-that were mostly expressed by the M cell populations identified in both murine tissues (**Figures 6B, C**). These four DEGs also were expressed in other epithelial cell populations in murine lungs (**Figure 6B**), while they were almost exclusive to M cells in mouse trachea (**Figure 6C**). Other human adenoid mature M cell DEGs like *CEMIP*, *MYBPC1*, *PPCDC*, *ENPP2*, *XKR4*, and *TMEM178B* had very low or absent expression in murine airway M cells as well as in the other epithelial cell types. In the case of human adenoid immature M cell DEGs, no consistent pattern emerged from either mouse atlas.

We also compared our findings with a population of M cells from the GI tract, specifically M cells identified in mouse gut organoids treated with RANKL ^72^ (**Figure 6D**). This *in vitro*-induced M cell population showed significant expression of the mature M cell DEGs *FRK*, *ARHGAP26*, *PARD3B*, and *INSR*, similar to mouse lung and tracheal M cells (**Figures 6B, C**). We also found that the adenoid immature M cell markers *DOCK8*, *PLGC2*, and *TNFAIP8* were expressed by the RANKL-induced GI M cell population.

Taken together, our detailed comparative analysis demonstrated that human adenoid M cells express a common set of genes not only with M cells residing in other human MALT tissue like the palatine tonsil, but also with M cells in the murine airway and M cells differentiated *in vitro* by RANKL treatment of mouse GI organoids. Notably, we identified genes uniquely expressed by human adenoid M cells, emphasizing the impact of spatial localization on the M cell transcriptome.

## Discussion

In this work, using single cell nuclear transcriptomics, we provide a detailed annotation of both immune and non-immune cells within the human adenoid. Furthermore, by analyzing the data using bioinformatic tools, we identified several unexpected cell types and states in the human adenoid, including HSCs, an unknown epithelial cell type with a robust interferon signature and immature and mature M cells. We also used trajectory analysis to identify the stepwise maturation of cells within the adenoid.

Immune cells in the adenoid may share a common progenitor stem cell. Such stem cells are typically found in the bone marrow and peripheral blood, but other authors have described the presence of adult stem cells with multi-lineage differentiation capacity in secondary lymphoid organs like the tonsils ^104,105^, and human lung ^106^. Our identification of a common precursor for all the immune cell types in the adenoid may also help clarify the ontogeny of tissue-resident follicular DCs, as they previously have been assigned to a mesenchymal origin in human lymphoid follicles ^31–33^.

Although it was not the main goal of this study, we were able to identify all the principle epithelial cell types in the human adenoid including neuroendocrine cells that were recently reported to be absent from other human nasal mucosa locations like the lower nasal turbinates ^107^. Also, previous publications in nasal tissue were not able to distinguish some terminally differentiated cell types from their lineage originating cell cluster because of their shared features and the lack of expression of established cell markers ^50^, highlighting the importance of using alternative tissues like the adenoid to identify and trace upper airway cell lineages. One notable cell type that we did not identify in the adenoid was the pulmonary ionocyte that were originally described in medium and lower airways in mice ^47,48^. This observation may reflect species differences, an inability to capture these nuclei with the technology used in our study, or simply that ionocytes develop lower in the respiratory tract. Our work, combined with the work of others, paves the way to better characterize the different epithelial cell types and their roles in mucosal biology.

We identified a cell type in the subset analysis of the basal/hillock lineage cluster with a unique interferon-related gene signature that we named UCT cells. This cluster, though seemingly derived from basal cells, lacked similarity with other known cell types in the human adenoid. The transcriptional profile of UCT cells was consistent with detection of and response to nucleic acids and induction of interferon and pro-inflammatory cytokines. Further analysis will be required to confirm the presence and functional significance of the UCT cell in the adenoid.

In our subset analysis of the club cell lineage, we identified two distinct populations of M cells in the human adenoid. Both M cell populations showed variable expression of traditional M cell markers previously used to define gut M cells. It may be that differences in techniques, namely snRNA-seq presented here versus immunofluorescence and flow cytometry, explain the discrepant results. Recently, CR2 and CORO1A have been defined as uniquely expressed in mouse lung and trachea and human Peyer’s patches M cells respectively ^17,18,108^. However, in the human adenoid M cell populations, expression of both genes was very low and was also not unique to M cells as other cell clusters showed higher expression levels. Likewise, expression of the adenoid immature and mature M cell DEGs in other datasets was variable. Notably, while some genes were unique to human adenoid M cells, a set of 4 genes including *INSR*, *PARD3B*, *ARHGAP26* and *FRK* identified from human adenoids were also expressed in mouse tracheal, lung and organoid samples, suggesting that they may represent a distinct M cell signature across species.

Differences in gene expression between airway and GI tract M cells may be a consequence of the distinct environments and conditions to which both populations of M cells are exposed in mouse and human tissues, underscoring the significance of their specific locations within the human body ^22,107^. For example, in the GI tract, gut-innervating neurons regulate both M cell function and segmentous filamentous bacteria (SFB) growth via production of calcitonin gene-related peptide ^109^. Also, hypoxia has been recently identified as key for the induction of epithelial cell differentiation in human lungs ^110^. Whether factors like the specific neurons innervating the human adenoid and the unique microbiome of the upper airway/oropharynx compared to the GI tract, or exposure to different oxygen and carbon dioxide concentrations impact adenoid M cell development in a similar or distinct manner to GI M cells is unknown. There are limitations to our study. First, we found variability in cell clusters among the donor samples. Such variability may be due to biologic variation or potentially to the kits used in library preparations and sequencing as has been previously reported ^111,112^. Second, because these tissues were removed from children undergoing elective adenoidectomy, it is not possible to definitively exclude the possibility that some transcripts were abundant owing to asymptomatic infection. For example, though we identified the UCT as distinct from other cells, whether their profile reflects a cell population poised to respond to infection in the upper airway or a cell type that was already infected at the time of organ collection cannot be determined by this analysis. Finally, snRNA-seq only analyzes nuclear transcripts and has previously shown difficulties quantifying transcripts with a low nuclear abundance ^113^. Consequently, some studies have reported different cell proportions when snRNA-seq and single-cell RNA sequencing (scRNA-seq) have been directly compared on the same sample ^114–116^.

Elucidation of the complete cell repertoire in the human adenoid will facilitate studies aimed at dissecting the earliest interactions of airway pathogens and the mucosal immune system. Furthermore, characterization of human adenoid M cells, which may share features with M cells in the lower respiratory tract ^17,18^, enhances and deepens our understanding of initial mucosal immune responses and functions. Such knowledge could generate urgently needed new approaches for infectious disease treatment and vaccine development at the earliest contact points ^117^.

## Materials and methods

### Sample collection

Samples were collected from 6 donors (age 2 – 14; 3 males and 3 females) undergoing elective adenoidectomy for obstructive sleep apnea. All samples were obtained at the Children’s Medical Center of Dallas. All donors or legally authorized representatives gave informed consent, which was approved by the University of Texas Southwestern Institutional Review Board (IRB #STU062016-087).

### Generation of single-nucleus suspensions

Isolation of single nuclei from human adenoid tissue was carried out by adapting a previously described protocol ^118^. Briefly, excised adenoids were immediately placed on ice after collection. Adenoid tissue was then placed in chilled homogenization buffer (0.25 M sucrose, 150 mM KCl, 5 mM MgCl_2_, 0.1 M Tris buffer pH 8.0, 0.1 mM DTT, 0.1 % Triton-X, and 0.2 U/µl RNase inhibitor) and minced. Samples were individually transferred to a gentleMACS C tube and homogenized with a gentleMACS dissociator. The tissue homogenate was passed through a 70 µm filter and diluted 1:1 with a chilled homogenization buffer. After centrifugation at 500g for 5 minutes at 4°C, nuclei were resuspended in wash buffer (1% BSA in DPBS with 0.2U/µl RNase inhibitor). Nuclear integrity was then assessed by microscopy, and the number of nuclei was counted using a BioRad TC20 automated counter. Only clump- and debris-free suspensions with at least 75-90% of nuclei with intact and smooth nuclear membranes were used for sequencing.

### Nuclear sequencing

Nucleus suspension from each donor were fixed using the Parse Evercode Nuclei Fixation v1 (samples 1 – 4) or v2 (samples 5 – 6). Since not all the samples were collected on the same day, all samples were processed until fixation first. Two independent libraries were prepared using the Parse Evercode Whole Transcriptome kit (V1 or V2) and sequenced on an Illumina NextSeq 2000.

### Sequencing data analysis and cell type scoring

Sequencing data were processed, mapped to the human (GRCh38) genome, and demultiplexed using Parse pipeline v1.1.2. The resulting matrix was normalized and scaled using Seurat R package (version 4.3.0) standard pipeline^119^ in R version 4.2.3^120^. Since adenoid samples were prepared with different kits, individualized quality control thresholds were used based on genes per nuclei and percent mitochondrial RNA reads following Parse recommendations. Briefly, for adenoids 1 and 2, we excluded nucleus barcodes with <150 and >2,000 detected genes and a mitochondrial expression >8 or 4.5% respectively; for adenoids 3 and 4, <100 and >2,000 detected genes and >20% mitochondrial counts; and for adenoids 5 and 6, <200 and >2,000 detected genes and >2.3 or 0.24% respectively of mitochondrial counts. A shared embedding was calculated for sample integration using Harmony^121^ before clustering and Uniform Manifold Approximation and Projection (UMAP) plot was used to visualize the clusters^122^. Only positive markers, expressed in at least 25% of the nucleus population with a log2 Fold-Change equal to or higher than 0.25, were identified for each cluster using the FindMarkers and FindAllMarkers tests integrated in Seurat. The identity of nucleus clusters was determined by manually cross-referencing top differentially expressed genes with previously published studies. Once the epithelial-related clusters were identified, these clusters were subset and re-clustered following the same procedure explained above. The subcluster annotation was performed in the same way as the parental population.

### Trajectory analysis

Single-nucleus trajectories were built with the software Monocle 3 (version 1.3.5)^53,54^ for individual epithelial cell subclusters. Specifically, single-cell trajectories were performed for the 2,796 nuclei from basal/hillock lineage or the 2,469 nuclei from the club lineage. Root nodes were selected based on previous knowledge about the precursor cell type for each analyzed lineage –basal and hillock in basal/hillock lineage and club in club lineage–, and by manually choosing a midpoint of these cell populations. After this, cells were colored and ordered in boxplots by pseudotime values to determine the degree of cell differentiation for each individual nucleus. Finally, genes that changed as a function of pseudotime were analyzed and plotted in the corresponding trajectory graph. The percentages shown in each trajectory graph represent the correlation of each individual gene with a trajectory pathway.

### Pathway enrichment analysis

Functional annotation of the differentially expressed genes for a specific cell cluster was performed with the Database for Annotation, Visualization, and Integrated Discovery (DAVID) tool ^55,56^. The list of genes specified in the corresponding text sections was used to study the most relevant biological processes of the unknown cell type and mature M cells. Only biological processes with a q-value lower than .05 were considered for each cell type.

### Ligand-receptor analysis

To study intercellular communication, metadata and data slots of the Seurat object were used to generate a CellChat object using the CellChat R package (CellChat 1.6.1) ^57^. Data were preprocessed using standard CellChat workflow. Human CellChat database of known ligand-receptor interactions was used to identify the connections between all cell types identified in the human adenoid.

### Human adenoid immunofluorescence

Human adenoid sections were obtained as previously described ^20^. Briefly, samples were fixed with 10% neutral buffered formalin solution (Sigma-Aldrich), embedded in paraffin, sectioned (5 µm), and mounted on glass slides. Slides were deparaffinized using xylene and ethanol washes followed by heat mediated antigen retrieval in 10mM sodium citrate (pH 6.0). After this, tissue sections were permeabilized and blocked in 0.4% Triton X-100 and 5% bovine serum albumin (BSA) in PBS (blocking solution with Triton) for 1 hour at room temperature. Slides were washed with PBS and incubated with a 1:100 dilution of rabbit anti-TNFAIP2-CF647 (Biorbyt #orb101923-CF647), rat anti-NKM 16-2-4-PE (Miltenyi #130-102-150), and mouse anti-CEMIP (KIAA1199)-FITC (Bio-Techne #NBP2-50336F) in blocking solution with Triton overnight at room temperature. Slides were then washed with PBS, incubated with DAPI, washed with PBS, mounted in Prolong Gold antifade reagent, and imaged using a CSU W1 spinning disk confocal microscope (Nikon).

## Acknowledgements

The authors thank Joe Musmacker, Brian Walsh, Daniel Diaz, and Cooper Siepmann from Parse Biosciences for providing technical and computational support. The authors also thank core facilities at UT Southwestern Medical Center for their important contributions to this work including the UT Southwestern McDermott Center Next Generation Sequencing (NGS) Core, particularly Caitlin Eaton and Vanesa Schmid; the Histopathology Core, particularly the core director Bret Evers and John Shelton; and the Quantitative Light Microscopy Core, a Shared Resource of the Harold C. Simmons Cancer Center, supported in part by an NCI Cancer Center Support Grant, 1P30 CA142543-01.

## Data availability

All snRNA-seq data have been deposited in Gene Expression Omnibus (GEO) and will be available at the time of publication.

## Author contributions

Conceptualization, S.A.A., M.U.S.; Formal analysis, S.A.A., K.M., A.W., J.T.L., M.U.S.; Computational analysis, S.A.A., K.M., A.W., J.T.L.; Sample collection, R.B.M.; Investigation, S.A.A., G.G., I.S.; Funding acquisition, T.J.P., M.U.S.; Project administration, M.U.S.; Supervision, M.U.S.; Writing – original draft, S.A.A, M.U.S.; Writing – review and editing, All authors.

