## Supplemental Figures and Legends for "Single cell transcriptional analysis of human adenoids identifies molecular features of airway microfold cells"

### Supplemental Figure 1

**A**

| Cell type | Marker gene | References |
|---|---|---|
| Hemat. stem cells | PTPRC | 37 |
| | ATXN1 | 38 |
| | CD38 | 39 |
| | SLAMF1 | 40 |
| B cells | CD19 | 41 |
| | PXK | 42 |
| | CD74 | 43 |
| T cells | CD247 | 44 |
| | TRBC2 | 45 |
| | CD3E | 46 |
| Dendritic cells/Macs | TFEC | 47 |
| | ITGAX | 48 |
| | ZBTB46 | 49 |
| Follicular DCs | CR1 | 50 |
| | CR2 | 50,51 |
| | CXCL13 | 50 |
| Basal/hillock lineage | IGFPB3 | 52 |
| | KRT8 | 53 |
| | PSMD5 | 53 |
| Club lineage | CDHR3 | 53 |
| | FOXJ1 | 53 |
| | MUC5B | 53 |

**B**

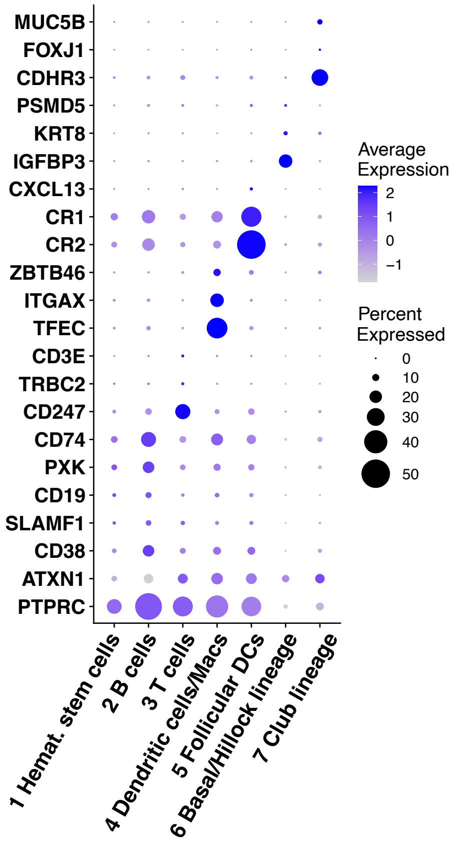


**C**

| Cell type | Marker gene | References |
|---|---|---|
| Basal cells | IGFPB3 | 52 |
| | DLK2 | 60 |
| | LAMB3 | 61 |
| Neuroendocrine cells | PSMD5 | 53 |
| | NEB | 60 |
| | PEX5L | 18 |
| Tuft cells | ANXA4 | 60 |
| | SPIB | 53 |
| | SOX9 | 53 |
| Unknown cell type | | |
| Hillock cells | KRT4 | 53 |
| | SCEL | 60 |
| | SPRR1B | 60 |

**D**

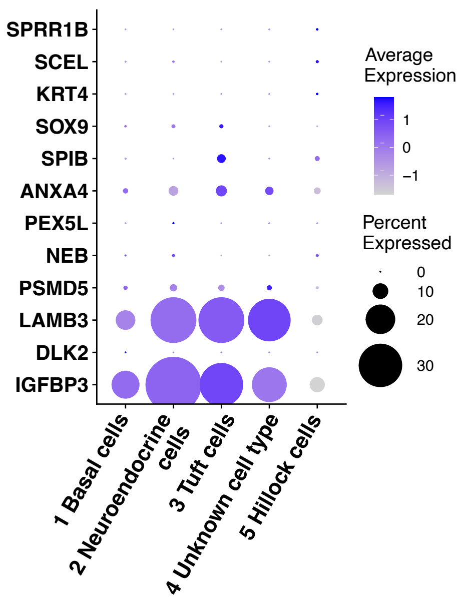


**E**

| Cell type | Marker gene | Reference |
|---|---|---|
| Club cells | SCGB1A1 | 53,60 |
| | NFIA | 53 |
| Ciliated cells | CDHR3 | 53 |
| | FOXJ1 | 53 |
| | SNTN | 60 |
| Goblet-1 cells | MUC5AC | 53 |
| | MUC5B | 53 |
| Goblet-2 cells | MUC4 | 53 |
| | IL19 | 70 |
| | CSF3 | 70 |
| Immature M cells | RELA | 13 |
| | RELB | 13 |
| | SOX8 | 13 |
| Mature M cells | SPIB | 59 |
| | TNFRSF11A | 59 |
| | TNFAIP2 | 75 |

**F**

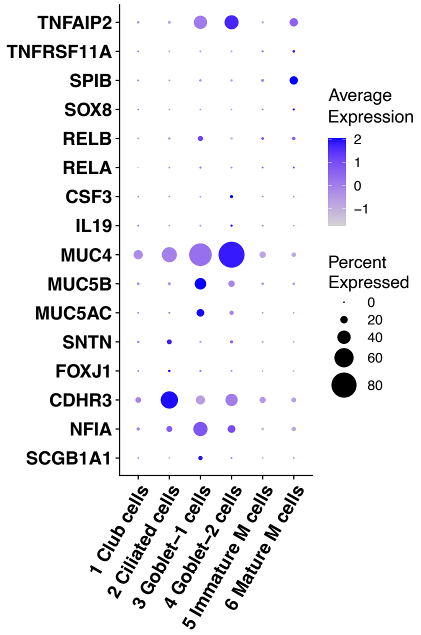

#### Supplemental Figure 2

A

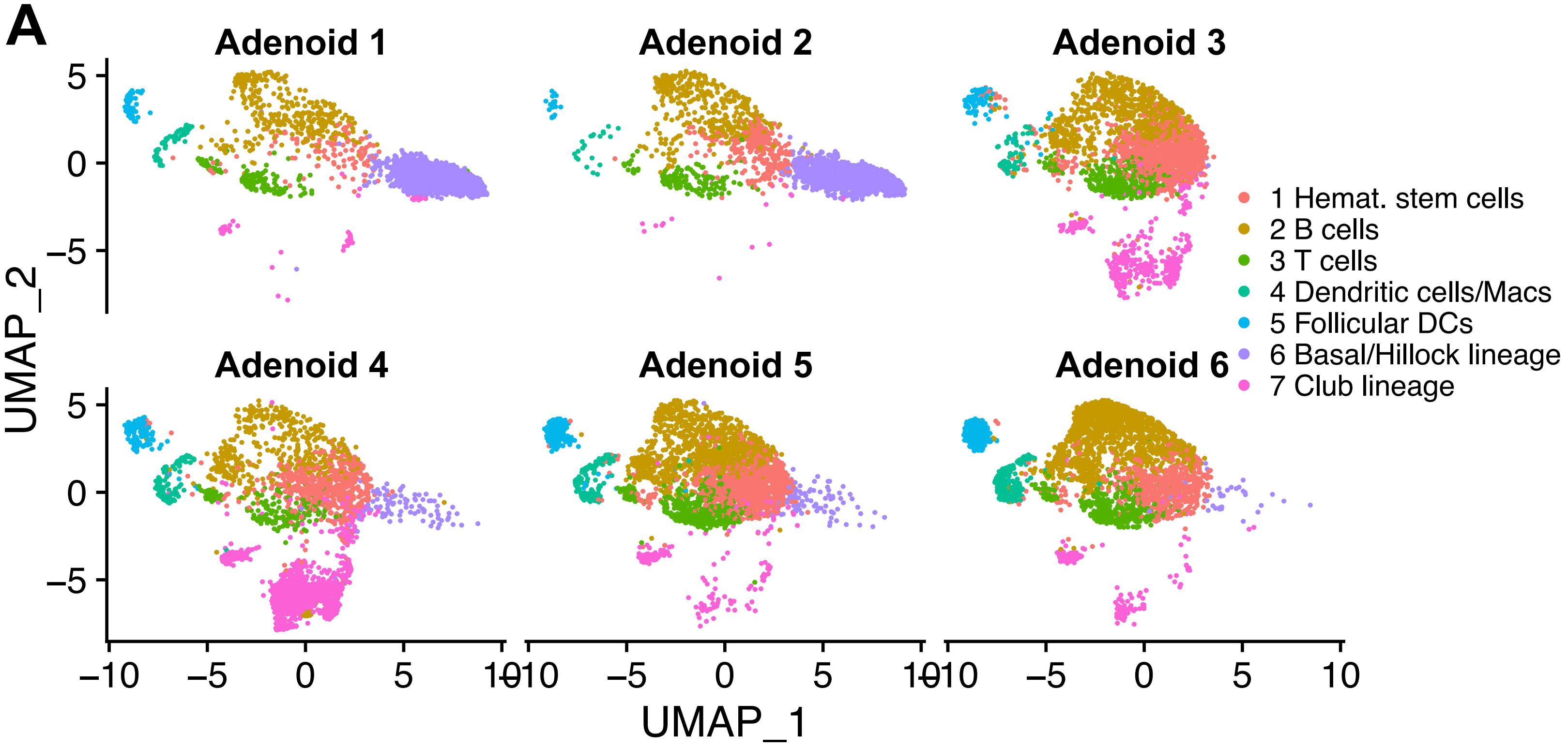


B

| | # nuclei | median_nCount | median_nFeat |
|---|---|---|---|
| **Adenoid 1** | 2,030 | 343 | 297 |
| **Adenoid 2** | 2,183 | 302 | 267 |
| **Adenoid 3** | 2,906 | 212 | 185 |
| **Adenoid 4** | 3,208 | 184 | 156 |
| **Adenoid 5** | 3,192 | 340 | 300 |
| **Adenoid 6** | 3,260 | 337 | 300 |

C

| | # nuclei | median_nCount | median_nFeat |
|---|---|---|---|
| **Hemat. stem cells** | 3,995 | 235 | 212 |
| **B cells** | 4,344 | 456 | 391 |
| **T cells** | 1,700 | 306 | 268 |
| **Dendritic cells/Macs** | 706 | 434 | 378 |
| **Follicular DCs** | 769 | 388 | 343 |
| **Basal/Hillock lineage** | 2,796 | 254 | 224 |
| **Club lineage** | 2,469 | 224 | 189 |

D

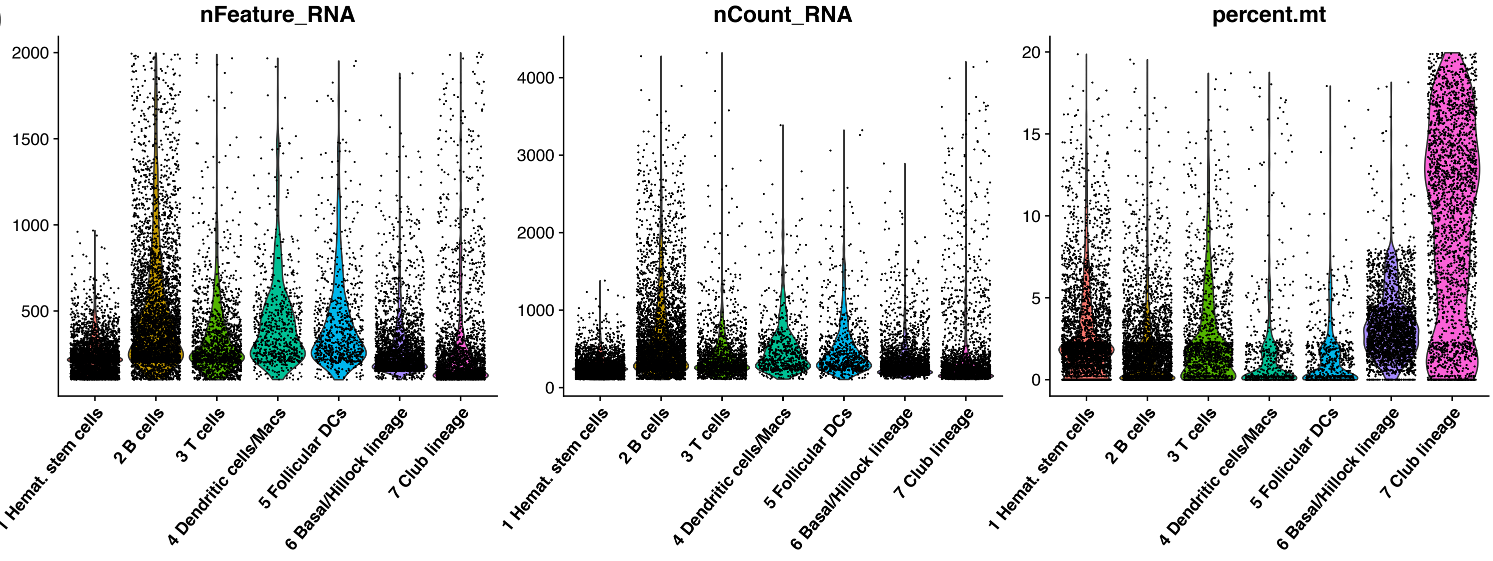

#### Supplemental Figure 3

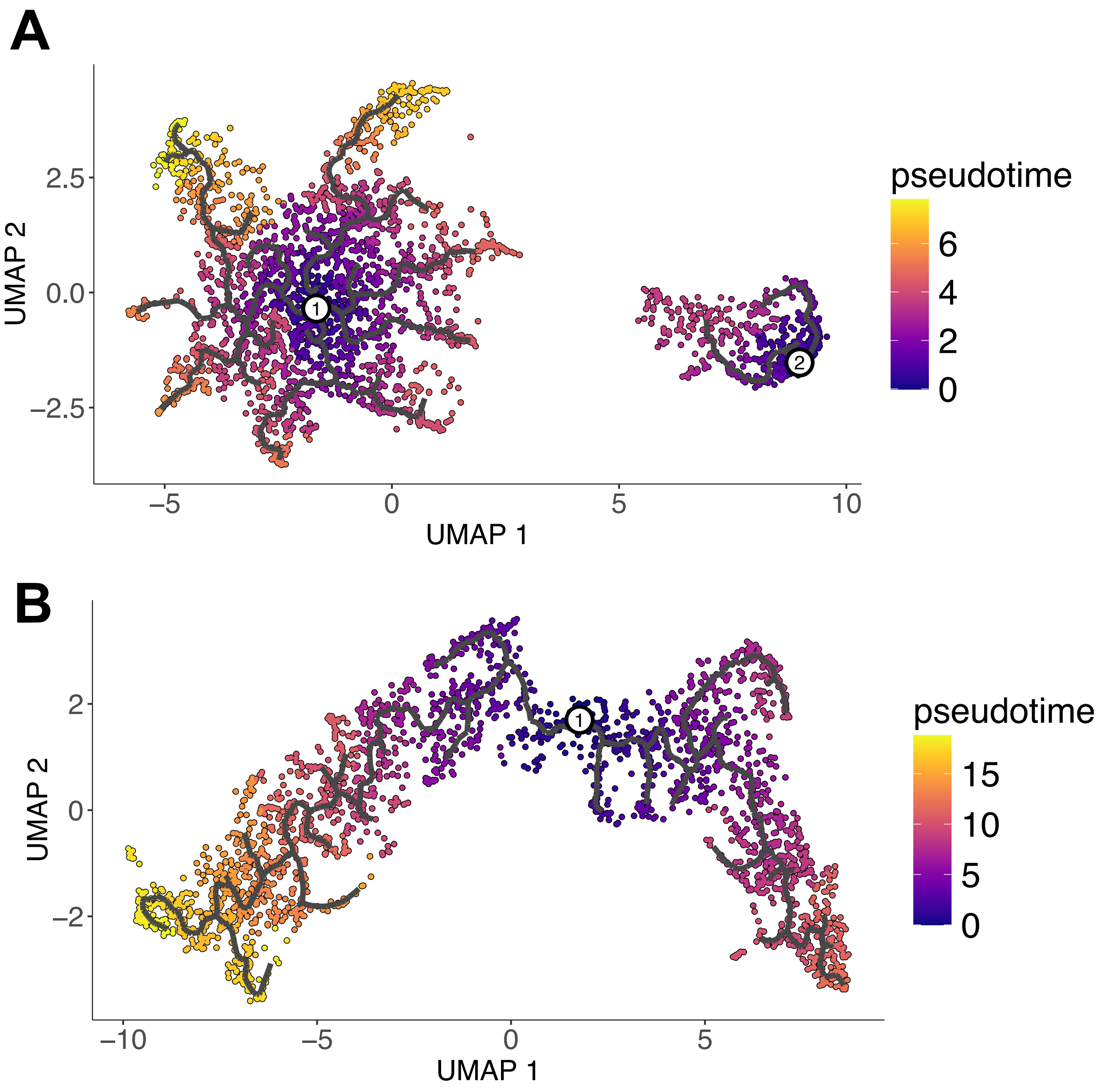

### Supplemental Figure 4

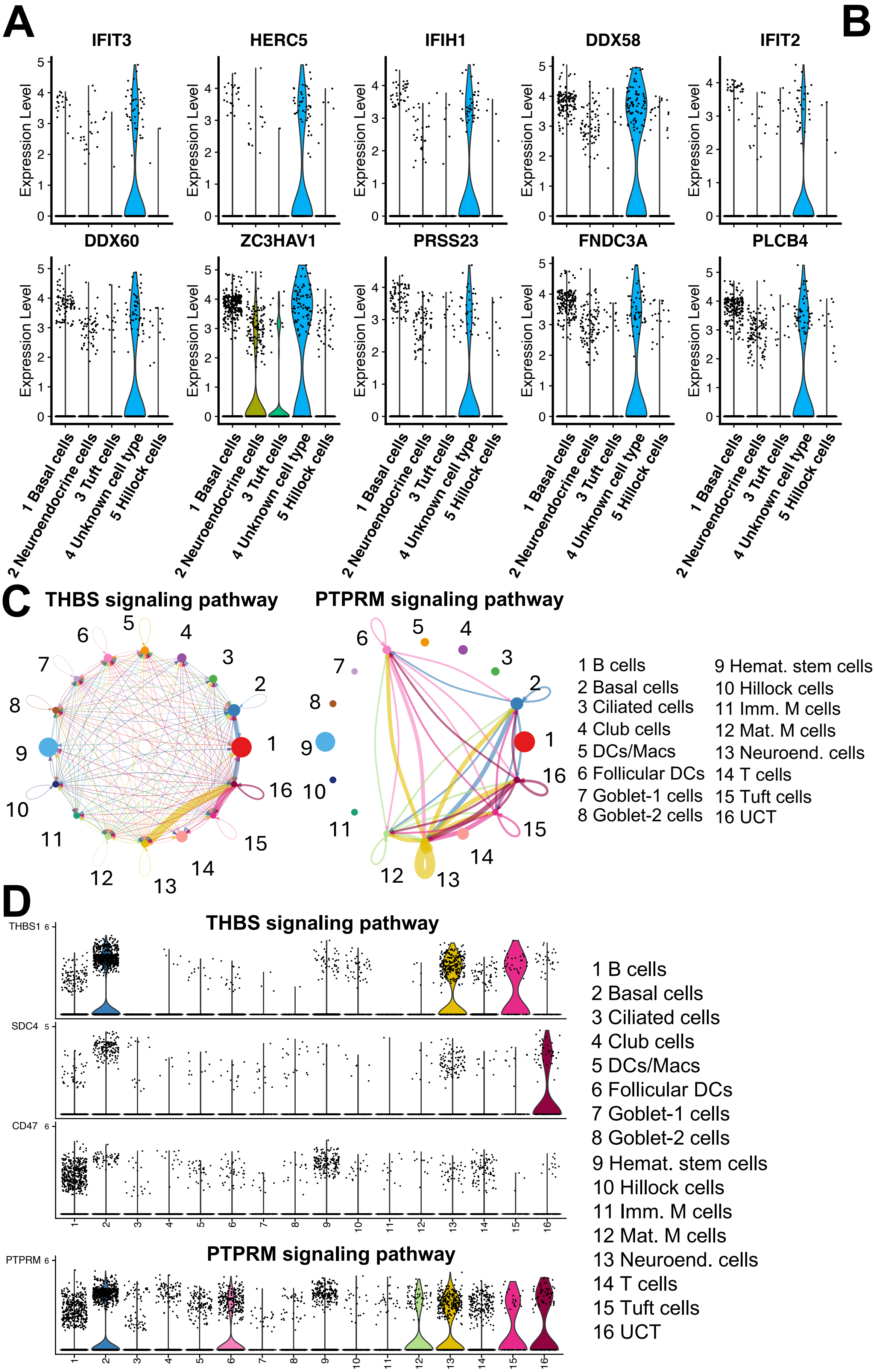


**B**

| Basal/hillock lineage only | Common | All populations |
|---|---|---|
| LUC7L3 | HERC5 | PHLDB2 |
| RBFOX2 | IFIT3 | MT-RNR1 |
| | DDX58 | DST |
| | IFIH1 | MIR924HG |
| | IFIT2 | HIF1A-AS3 |
| | ZC3HAV1 | CFTR |
| | DDX60 | DIAPH3 |
| | PRSS23 | YAP1 |
| | FNDC3A | RBFOX2 |
| | PLCB4 | EGFR |
| | SDC4 | STK3 |
| | KDM6A | TANC2 |
| | SPAG9 | AFDN |
| | MAP4K4 | AHNAK |
| | B4GALT5 | FNDC3B |
| | PARP14 | TRIO |
| | MAST4 | PTK2 |
| | ANKRD10 | EZR |
| | NFAT5 | HUWE1 |
| | SH3KBP1 | MALAT1 |
| | FRYL | IMMP2L |
| | PICALM | MYO1E |
| | EXT1 | RAD51B |
| | LAMB3 | RUNX1 |
| | PTPRM | EXOC4 |
| | FN1 | ZFAND3 |
| | LAMA3 | JMJD1C |
| | MACF1 | WWOX |
| | ATP11A | SYNE2 |
| | CTNND1 | |
| | NEAT1 | |
| | NF1 | |
| | SIPA1L1 | |
| | PPP6R3 | |
| | LRBA | |
| | PVT1 | |
| | HNRNPA2B1 | |
| | RNF213 | |

### Supplemental Figure 5

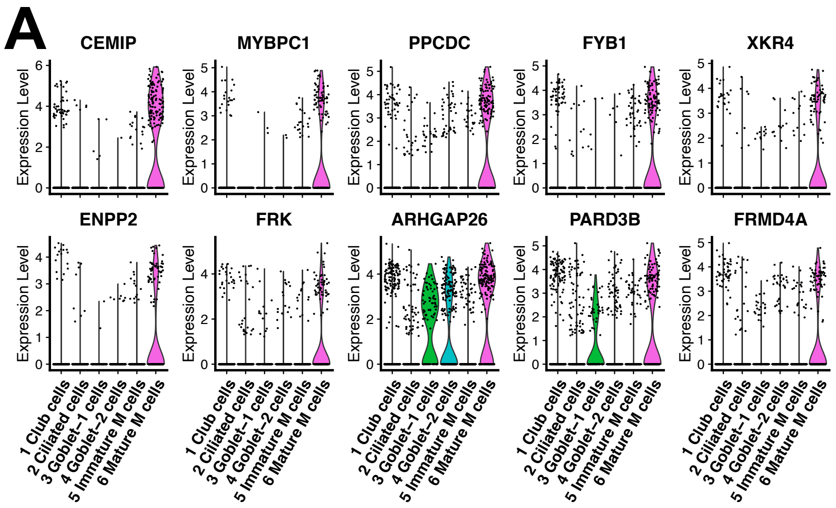


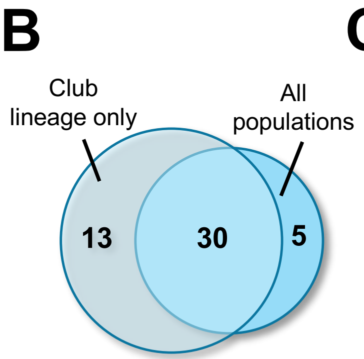


C

| Club lineage only | Common | All populations |
|---|---|---|
| MALAT1 | MYBPC1 | NEBL |
| ELMO1 | CEMIP | ANK3 |
| LPP | PPCDC | NEAT1 |
| ARHGAP15 | FRK | LDLRAD4 |
| TBC1D5 | ARHGAP26 | PDE4D |
| RBM6 | PARD3B | |
| SMYD3 | ENPP2 | |
| MACF1 | XKR4 | |
| FOXP1 | INSR | |
| PDE4D | TMEM178B | |
| TBC1D22A | AGAP1 | |
| AKAP13 | FYB1 | |
| MBNL1 | MAGI1 | |
| | MYO6 | |
| | FRMD4A | |
| | SDK1 | |
| | KIAA1217 | |
| | SLC9A9 | |
| | PTPRM | |
| | CTNND1 | |
| | ETV6 | |
| | ARL15 | |
| | FTX | |
| | 7SK | |
| | EML4 | |
| | PLEKHG1 | |
| | DDX5 | |
| | TRAPPC9 | |
| | JAZF1 | |
| | LYN | |

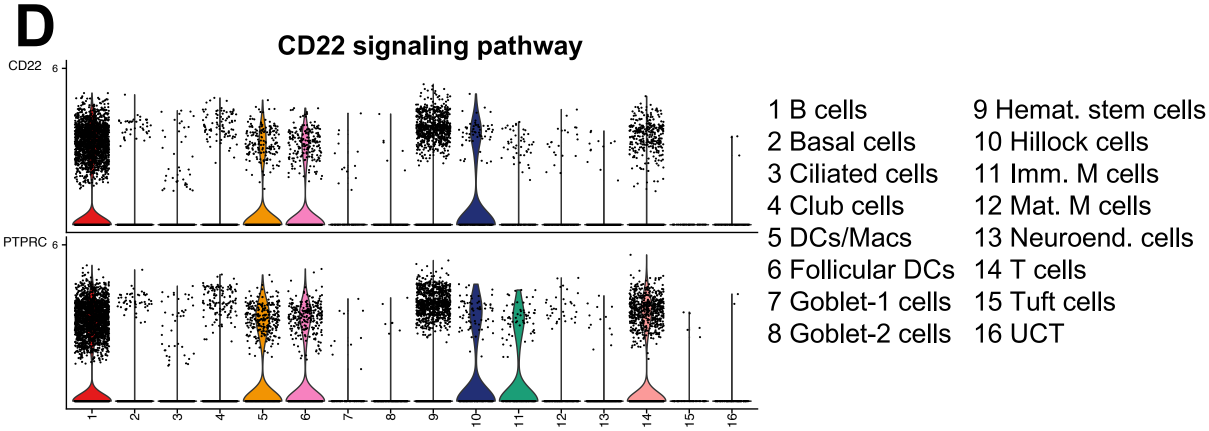

**Supplemental Figure 1. Cell marker genes used to identify each cell population in the human adenoid. Related to figures 1, 2, and 4.** Tables of markers used to identify cell populations in human adenoid (A), in basal/hillock lineage (C), and in club lineage (E). Only three representative markers for each cell population are shown. Dot plot of the average expression of cell population markers for human adenoid (B), basal/hillock lineage (D), and club lineage (F).

**Supplemental Figure 2. Cell composition of the human adenoid. Related to figure 1.** (A) UMAP plots of the 16,779 nuclei from human adenoids split by sample. Number of nuclei, median number of RNA fragments (median_nCount), and median number of genes in each nucleus (median_nFeat) detected in each human donor (C) or labeled nucleus cluster (D). (E) Violin plot of the number of genes detected (nFeature_RNA), number of RNA fragments (nCount_RNA), and percentage of mitochondrial related fragments (percent.mt) in each nucleus cluster of the human adenoid.

**Supplemental Figure 3. Pseudotime analysis of basal/hillock and club lineages. Related to figures 2 and 4.** Trajectory analysis of the basal/hillock lineage (A) and club lineage (B) showing the manually designated root nodes with the numbered label and colored by pseudotime. In the case of the basal/hillock lineage, one separated root node was designated for each trajectory group. Pseudotime color values are showed in the scale bar.

**Supplemental Figure 4. The "unknown cell type" is a basal cell progeny with a unique interferon-related gene signature. Related to figure 3.** (A) Individual expression levels in transcripts per million (TPM) of the top 10 UCT DEGs in the nucleus populations identified in the basal/hillock lineage only. (B) List of UCT DEGs present only when the basal/hillock lineage populations were used, when all the human adenoids populations were used, and common to both analyses. (C) Main signaling pathways in which UCT cells are involved either as a source or as a receptor. The numbers that correspond to each cluster are on the side. (D) Expression levels of the genes of the most significant ligand-receptor interactions in which UCT cells are involved.

**Supplemental Figure 5. Characterization of human adenoid M cells. Related to figure 5.** (A) Individual expression levels in transcripts per million (TPM) of the top 10 DEGs for mature M cells in club lineage only. (B) Venn diagram showing the overlap between the mature M cell DEGs identified only using club lineage populations or all the human adenoids ones. (C) List of mature M cell DEGs present only when the club lineage populations were used, when all the human adenoids populations were used, and common to both analyses. (D) Expression levels of the genes of the most significant ligand-receptor interactions in which mature M cells are involved.
